# MYC plus class IIa HDAC inhibition potentiates mitochondrial dysfunction in non-small cell lung cancer

**DOI:** 10.1101/2024.09.04.610116

**Authors:** Jina Park, Ying-Yu Chen, Jennie J. Cao, Julia An, Ray-Whay Chiu Yen, John D. Outen, Stephen B. Baylin, Michael J. Topper

**Author notes:** **Corresponding Author:** Michael J. Topper; The Sidney Kimmel Comprehensive Cancer Center, Johns Hopkins University School of Medicine, 1650 Orleans Street, Baltimore, MD 21287.

## Abstract

MYC is frequently activated in cancer, leading to significant efforts to develop MYC inhibitors. While much progress has been made in targeting MYC, combination treatment strategies are needed to exploit this molecular vulnerability. To this end, we interrogated transcriptome data from cancer cell lines treated with MYC inhibitors and identified HDAC5 and HDAC9, both class IIa HDACs, as therapeutic targets to inhibit concurrently. Notably, these HDAC isoforms, which can be specifically targeted by small molecules, are known augmenters of several hallmarks of cancer. The combination of MYC and class IIa HDAC inhibition induces a significant reduction in viability for NSCLC cell lines with high MYC and mitochondrial pathway activation. Additionally, combination treatment induces a robust reduction of MYC with concomitant elevation of mitochondrial ROS, both of which have a causal relationship with therapeutic efficacy. Confirmation of in vivo efficacy was pursued in several animal model systems, with subsequent molecular correlate derivation confirming the importance of MYC depletion and mitochondrial dysfunction in driving drug efficacy. Ultimately, we define a therapeutic approach combining MYCi and class IIa HDACi to potentiate anti-tumor efficacy in NSCLC.

## Introduction

Lung cancer is the leading cause of cancer-related deaths in both men and women, with non-small cell lung cancer (NSCLC) accounting for approximately 80-85% of all lung cancer cases.(1, 2) The primary factor contributing to the high mortality of NSCLC is advanced disease presentation at diagnosis, with 70-80% of patients harboring non-localized disease.(1, 3) Thus, the 5-year relative survival rate for NSCLC remains low at 26-47% and 6-11% for regional- and distant-stage cancer, respectively.(3) Currently, standard-of-care treatments for advanced NSCLC include surgery, chemotherapy, radiation, targeted therapy, immunotherapy, or combination multimodality treatment. However, achieving durable clinical benefit from standard therapies has been challenging and, in most cases, limited to only a minority of patients.(4–14) Identifying molecular vulnerabilities driving therapeutic response is critical for selecting patient populations most likely to benefit from a given therapeutic intervention. For example, somatic mutations in STK11/LKB1, which are prevalent in lung adenocarcinoma, especially among KRAS mutant tumors, have been reported to be associated with unfavorable outcomes in patients treated with broad classes of therapies, including immune checkpoint inhibitors, chemotherapies, and targeted therapies.(15–21) Thus, novel combination strategies to target NSCLC resistant to established therapeutic regimens represent an urgent clinical need.

The MYC oncoprotein is a pleiotropic transcription factor controlling the expression of genes promoting multiple hallmarks of cancer, including proliferation, cell survival, metabolism, genomic instability, angiogenesis, invasion, and immune evasion.(22–25) Aberrant regulation of MYC occurs in multiple cancer types, including NSCLC, which exhibits frequent MYC overexpression.(17, 26–32) Moreover, preclinical cancer models have demonstrated that MYC inactivation can elicit tumor regression, most notably in Kras-driven lung cancer models.(33–37) MYC has been proposed as a promising cancer therapeutic target, especially since side effects from MYC suppression are both tolerated and reversible, thus suggesting that a potential therapeutic window is achievable.(37–39) Pharmacological targeting of the MYC protein, however, has been challenging due to its intrinsically disordered structure and lack of an enzymatic active site.(40–42) Recently, two direct MYC inhibitors, Omomyc and MYCi975, have demonstrated improved in vivo efficacy and tolerability in preclinical studies.(38, 43) Both inhibitors suppress MYC function by impeding the interaction of MYC with its obligate binding partner, MAX. Omomyc, a dominant negative form of MYC, hinders the binding of MYC with MAX and target genes, inhibiting MYC-mediated transcription.(43) More recently, a small molecule MYC inhibitor, MYCi975, has been developed, which interferes with MYC-MAX binding and promotes the proteosome-mediated degradation of MYC.(38) Despite recent advances in targeting MYC, these agents have shown limited in vivo therapeutic efficacy as mono-therapies. Thus, combination treatments optimized to target additional molecular vulnerabilities are needed to leverage this novel approach to cancer treatment.

Histone deacetylases (HDACs) contain a highly conserved deacetylase domain, which removes acetyl groups from the lysine residues of histone and non-histone proteins.(44) These proteins can be broadly classified into 5 protein families, class I, IIa, IIb, III, and IV. HDAC overexpression is widely implicated in diverse cancer types, which makes HDAC inhibition a promising treatment approach.(44, 45) Particularly, the class IIa HDAC family, which includes HDAC4, HDAC5, HDAC7, and HDAC9, are emerging cancer targets due to their association with cancer progression, survival, and poor prognosis.(46–52) As opposed to class I HDACs, which are ubiquitously expressed, class IIa HDAC expression is both tissue-specific and elevated in lung cancer compared to adjacent normal tissue.(49, 50, 53, 54) Critically, the targeting of this HDAC family has not been extensively studied in contrast to pan-HDAC or class I HDAC-specific inhibition.

Here, we define a therapeutic paradigm wherein class IIa HDAC inhibitors are combined with MYC inhibition to potentiate treatment efficacy in NSCLC. Furthermore, through mechanistic studies, we reveal that combination treatment induces suppression of transcriptional programs involved in cell cycle progression and mitochondrial function. Additionally, we define a causal relationship between drug efficacy and both MYC depletion and oxidative stress. Finally, we translate our treatment paradigm to in vivo mouse models of NSCLC, including patient-derived xenograft and syngeneic models, and elucidate a conserved combination therapy-driven anti-tumor response, which is associated with a reduction in MYC, mitochondrial, and vascularization pathway activation.

## Results

### Combination of MYC and class IIa HDAC inhibition induces a substantial reduction of NSCLC cell line viability

To identify novel combinatorial drug strategies that potentiate MYC inhibition in cancer cells, we analyzed Gene Expression Omnibus transcriptomic data from cancer cell lines treated with either MYCi975 (GSE135800, PC3 prostate cancer cell line) or Omomyc (GSE126453, NSCLC cell lines).(38, 43) These data revealed 1189 and 3306 differentially expressed genes (DEGs) by MYCi975 and Omomyc treatment, respectively (Fig. S1, A and B). Among these DEGs, we specifically investigated HDACs, which are critical targets for cancer treatment via epigenetic therapy. MYC inhibition in both prostate and NSCLC cancer cell lines resulted in significantly augmented HDAC5 and HDAC9 expression, whereas expression of other HDAC isoforms was not significantly altered by treatment (Fig. S1, A and B). Interestingly, HDAC5 and HDAC9 are both class IIa HDACs, which have been proposed as promising cancer therapy targets. Critically, two class IIa HDAC-specific inhibitors, TMP269 and TMP195, have been recently identified as being both potent and highly selective.(55) We thus aimed to assess whether treatment efficacy might be enhanced by simultaneously targeting both MYC and class IIa HDACs in NSCLC. To evaluate this hypothesis, we tested drug sensitivity to the combination of MYCi975 plus TMP269 across a panel of 18 human NSCLC cell lines, which recapitulate the histological and mutational landscape of NSCLC. Interestingly, cell lines separated into two distinct groups based on area under the curve (AUC) analysis of the MYCi975 dose-response in the presence of 25 or 50 µM TMP269 (Fig. 1A). Therefore, we defined the group of cell lines with lower AUC as responders and those with higher AUC as non-responders. Importantly, these responding cell lines have higher combination drug sensitivity across multiple dose combinations tested compared to non-responders (Fig. 1, B and C, and Fig. S1, C and D). Furthermore, normalized dose-response curves demonstrated that in responders, TMP269 co-treatment resulted in a more substantial anti-proliferative effect than MYCi975 alone, whereas, in non-responders, there was no additional benefit from concurrent administration (Fig. 1, D and E, and Fig. S1, E and F). Based on these data, we next explored the relationship between the sensitivity to MYCi975 alone and the combination treatment by comparing the AUC values with or without TMP269 treatment in each cell line (Fig. 1F). This analysis revealed that the sensitivities to MYCi975 mono-treatment and the combination treatment are positively correlated (Fig. 1F). To explore whether a greater than additive effect was present, we performed synergy score calculation of dose-response data using the Bliss model.(56) This analysis identified higher synergy scores in responsive cell lines at concentrations with biologically meaningful drug effects than non-responders (Fig. 1, G and H, and Fig. S1, G and H). Additionally, drug antagonism at high concentrations of combination treatment was observed in 6 out of 8 non-responders but only 1 out of 10 responders (Fig. 1, G and H, and Fig. S1, G and H). To validate and extend these findings, we evaluated the combination of MYCi975 and another class IIa HDAC targeting agent, TMP195, which has demonstrated in vivo activity and anti-tumor efficacy.(57) Similar to the effects of the combination treatment with TMP269, MYCi975 plus TMP195 induced a substantial reduction in cell viability across four responsive cell lines (H460, H1703, H838, and H1755). In agreement with TMP269 dose-response data, combination TMP195 plus MYCi975 presented lesser treatment effects in two previously defined non-responsive cell lines (HCC827 and H441) (Fig. S2A). Furthermore, normalized dose-response curves and synergy calculation showed that synergistic or additive effects were also derived from MYCi975 plus TMP195 across multiple dose combinations in responsive cell lines (Fig. S2, B and C). We also demonstrated that the enzymatic activities of class IIa HDACs are significantly reduced upon both TMP195 and combination treatment (Fig. S2D). In summary, dual targeting of MYC and class IIa HDACs significantly decreased cell viability, with demonstrable synergy noted for most NSCLC cell lines tested.

**Figure 1.**
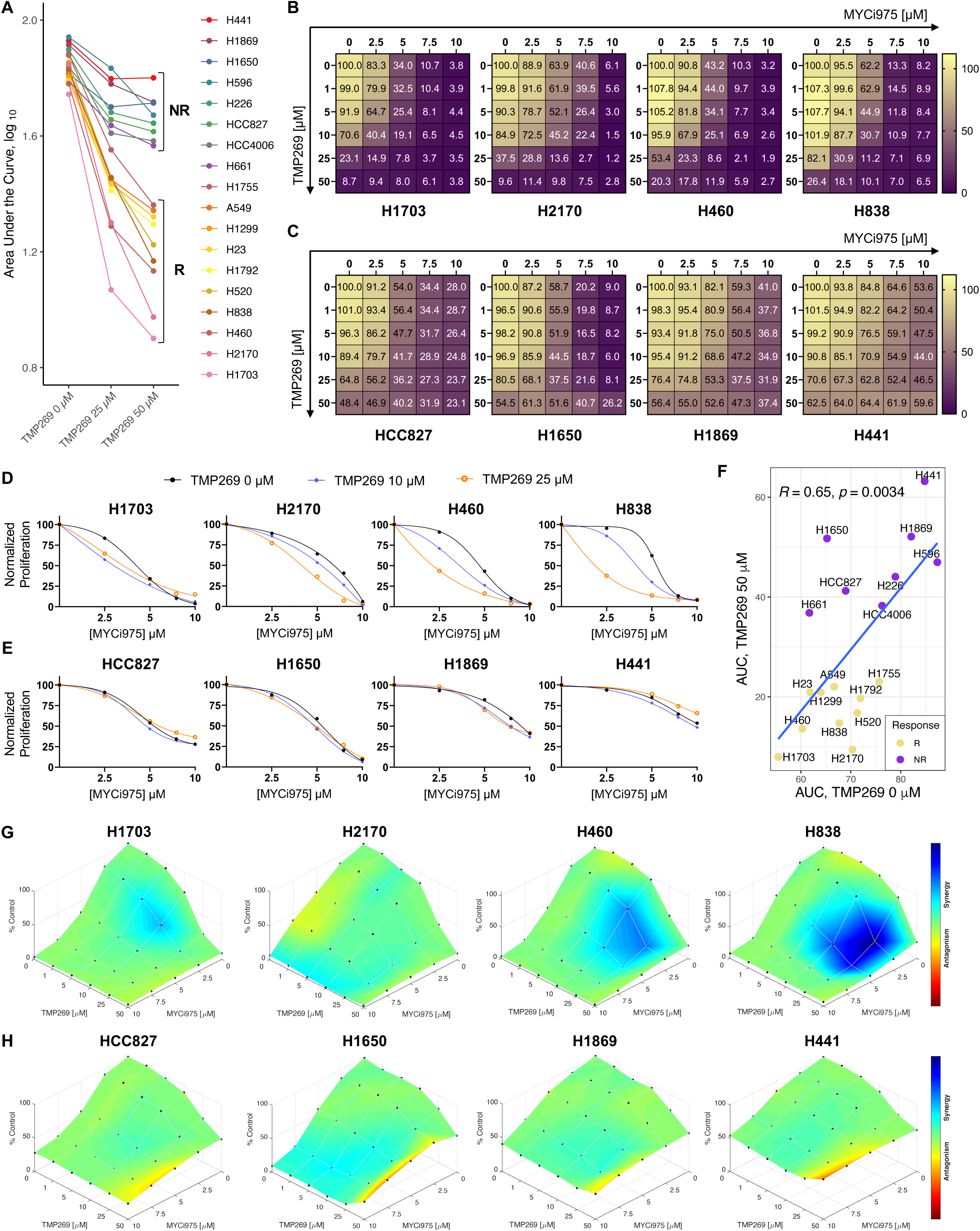
Treatment effects derived from the dual inhibition of MYC and class IIa HDACs in NSCLC cell lines. **A,** Area Under the Curve (AUC) of MYCi975 dose-response curves in the absence or presence of TMP269 (25, 50 µM). Each cell line was treated with MYCi975 (0, 2.5, 5, 7.5, 10 µM) +/− TMP269 (0, 25, 50 µM) for 72 h. R: responders, NR: non-responders. **B** and **C,** Dose-response matrices for percent viability in responsive **(B)** or non-responsive **(C)** cell lines. Values are reported as means from three independent experiments. Each cell line was treated with combination drugs as indicated for 72 h. Yellow indicates high viability, and purple indicates low viability. **D** and **E,** MYCi975 dose-response curves for normalized proliferation in the absence or presence of TMP269 (0, 10, 25 µM) in responsive **(D)** or non-responsive **(E)** cell lines. **F,** Scatter plot with AUC values for MYCi975 dose-response curves in the absence and presence of TMP269 50 µM for each cell line. Pearson correlation coefficient and p-value are depicted. **G** and **H,** Color on the dose-response surfaces represents the synergy score derived based on the Bliss model in responsive **(G)** or non-responsive **(H)** cell lines. Cool color indicates synergy and warm color indicates antagonism.

### MYC targets, reactive oxygen species, and mitochondrial pathways are basally activated in responsive cell lines compared to non-responders

Based on the distinct phenotypic separation of responders and non-responders, we sought to define baseline differences between these groups. To examine the differences in basal transcriptional profiles between responders and non-responders, we analyzed Cancer Cell Line Encyclopedia (CCLE) RNA-seq data. These data revealed 434 downregulated and 72 upregulated DEGs in responsive vs. non-responsive cell lines (Fig. S3, A and B).(58) Gene set enrichment analysis (GSEA) on RNA-seq data uncovered that responding cell lines have higher MYC targets, oxidative phosphorylation, and reactive oxygen species (ROS) pathway signatures relative to non-responding cell lines (Fig. 2A). Moreover, critical pathways in mitochondrial biology, such as mitochondrial central dogma, mitochondrial ribosome, and oxidative phosphorylation (OXPHOS), were upregulated in responsive cell lines compared to non-responsive cell lines, as identified by GSEA on MitoCarta 3.0 gene sets (Fig. 2B).(59) Since MYC target pathways are significantly enriched in responding cell lines; we next investigated MYC protein levels across our cell line panel using the CCLE proteomics dataset.(60) These data revealed higher normalized MYC protein levels in responders relative to non-responders (Fig. 2C). Interestingly, the derived AUC values, which are inversely associated with sensitivity, were negatively correlated with normalized MYC protein levels across the cell lines assayed (Fig. 2D). However, H1703, which is the most responsive cell line defined by AUC (Fig. 1A), had the lowest MYC protein levels among responders, thus suggesting that MYC levels while important, are not alone sufficient for driving the differential response to the combination treatment. Additionally, we noted no differences in normalized protein levels of class IIa HDACs between responders and non-responders (Fig. S3C). Next, we annotated the genomic alterations in genes with significantly recurrent mutations, copy number losses, or gains in NSCLC(17) for each cell line to associate these alterations with treatment efficacy (Fig. 2E). These data revealed that cell lines with EGFR genomic alterations were non-responders, whereas STK11 mutant cell lines were responders (Fig. 2E). Additionally, 6 out of 7 KRAS-altered cell lines were found to be responsive to the combination treatment. Fisher’s exact test found a statistically significant association between genomic perturbations in EGFR and KRAS (FDR-adjusted p-value < 0.1) and therapeutic response (Supplementary Table 1). Considering the varied patterns of MYC expression, as well as EGFR, STK11, and KRAS genomic alterations between responders and non-responders, we next investigated the relationship between MYC expression profiles and EGFR, STK11, and KRAS genomic perturbations in human primary tumor samples. Interestingly, in The Cancer Genome Atlas (TCGA) dataset, EGFR-altered lung adenocarcinoma (LUAD) tumors had significantly lower MYC expression than EGFR wild-type (Fig. S3D),(61) while KRAS-altered tumors had higher MYC expression than KRAS wild-type (Fig. S3E), which agrees with prior analyses.(62) STK11 mutant status was not correlated with MYC expression levels (Fig. S3F). These data show that therapeutic responses to combination treatment are associated with MYC, ROS, mitochondrial pathways, and EGFR, STK11, and RAS genomic alterations in NSCLC cell lines (Fig. S3K).

**Figure 2.**
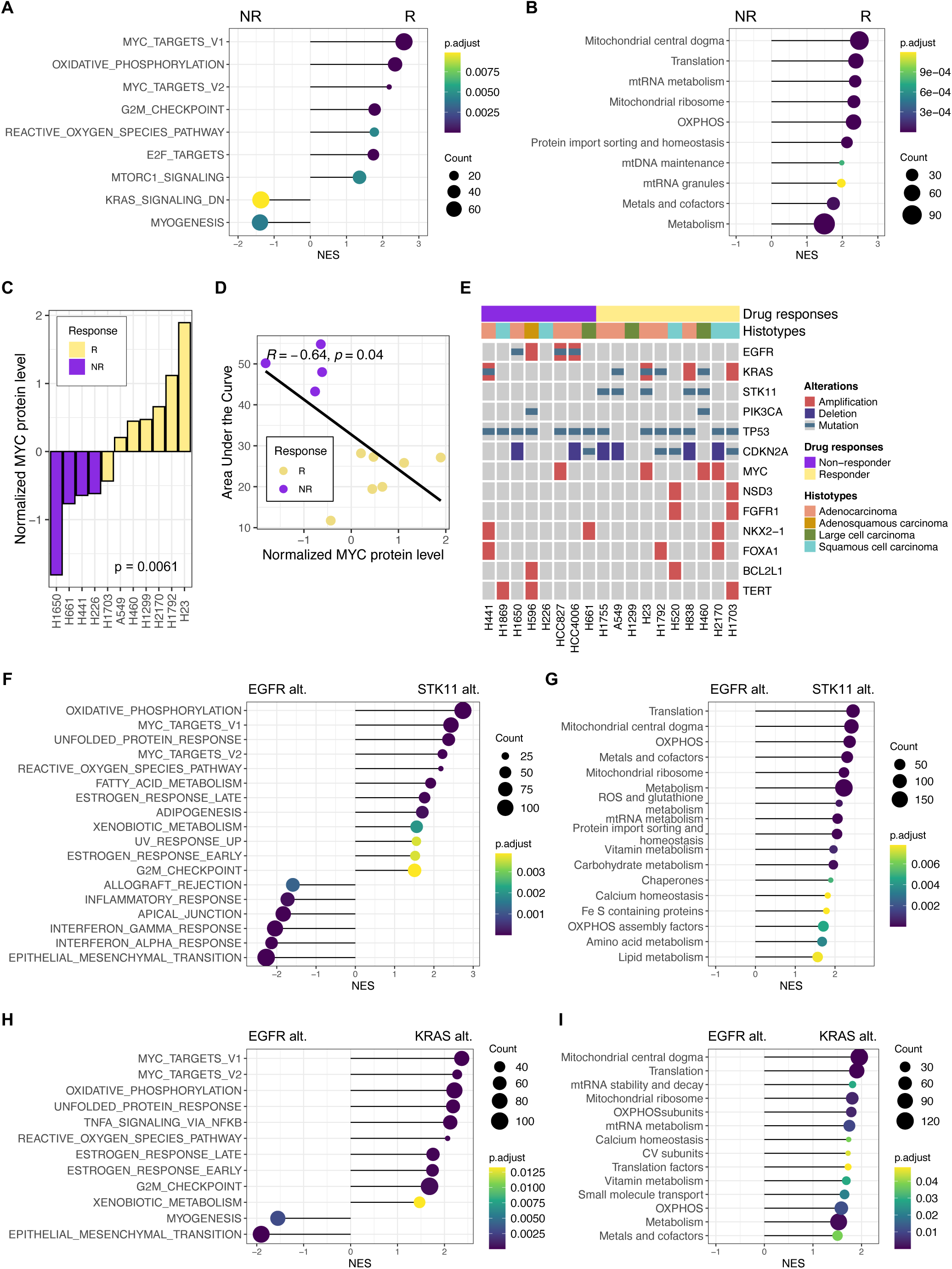
Transcriptional, genomic, and proteomic profiles associated with differential drug response. **A** and **B,** Gene Set Enrichment Analysis (GSEA) with Hallmark **(A)** or MitoCarta 3.0 **(B)** pathways based on differential expression analysis of responders (R) vs. non-responders (NR). Cancer Cell Line Encyclopedia (CCLE) RNA-seq data from the DepMap Portal was used for the analysis. Pathway dot plot, color indicates adjusted p-value, and size indicates gene count. **C,** Normalized MYC protein levels for 11 cell lines. Cell lines with missing values were excluded. CCLE proteomics data were obtained through the DepMap Portal. p-value was derived using the Mann-Whitney test. **D,** Scatter plot representing the correlation between the AUC values and the normalized MYC protein levels. The Spearman’s rank correlation coefficient and p-value are presented. **E,** Oncoprint depicting drug responses, histotypes, and genomic alterations in the driver genes of the cell lines. **F** and **G,** GSEA with Hallmark **(F)** or MitoCarta 3.0 **(G)** pathways based on differential expression analysis of STK11-altered vs. EGFR-altered TCGA LUAD samples. **H** and **I,** Gene Set Enrichment Analysis (GSEA) with Hallmark **(H)** or MitoCarta 3.0 **(I)** pathways based on differential expression analysis of KRAS-altered vs. EGFR-altered TCGA LUAD samples.

As expected based on previous reports, genomic alterations in EGFR and STK11 or KRAS were mutually exclusive in TCGA NSCLC samples (Fig. S3, G and H).(63–66) We next analyzed the TCGA LUAD RNA-seq dataset to identify transcriptional signatures associated with genomic alterations in these three genes. Principal component analysis (PCA) revealed that STK11-altered tumors are distinctively separated from tumors with EGFR genomic alterations (Fig. S3I), while the separation between KRAS- and EGFR-altered tumors was less profound (Fig. S3J). Pathway analysis on TCGA transcriptomic data found that tumors harboring STK11 or KRAS genomic alteration had higher MYC, ROS, and mitochondrial pathway enrichment when compared to the EGFR-altered tumors, recapitulating the pattern observed in CCLE responsive vs. non-responsive cell lines (Fig. 2, F to I).

### Combination of MYC and class IIa HDAC inhibition suppresses cell proliferation and induces apoptosis

Cell cycle assays using flow cytometry were performed in responsive cell lines to elucidate whether proliferation defects might drive the noted combination treatment effect. These assays revealed a combination treatment-induced significant increase in G0/G1 populations with an associated decrease in S phase, thus indicating that G1-S cell cycle arrest was resultant at 48 hours post-treatment initiation (Fig. 3, A and B). In H1703 specifically, G2/M phase populations were expanded by combination treatment, which is indicative of either G2-M cell cycle checkpoint induction or mitotic crisis (Fig. 3, A and B). Based on these cell cycle data, cell death was examined next. Longitudinal assessment of apoptosis revealed a significant increase in early and late apoptotic populations at 48 and 72 hours post-combination treatment initiation (Fig. 3, C and D).

**Figure 3.**
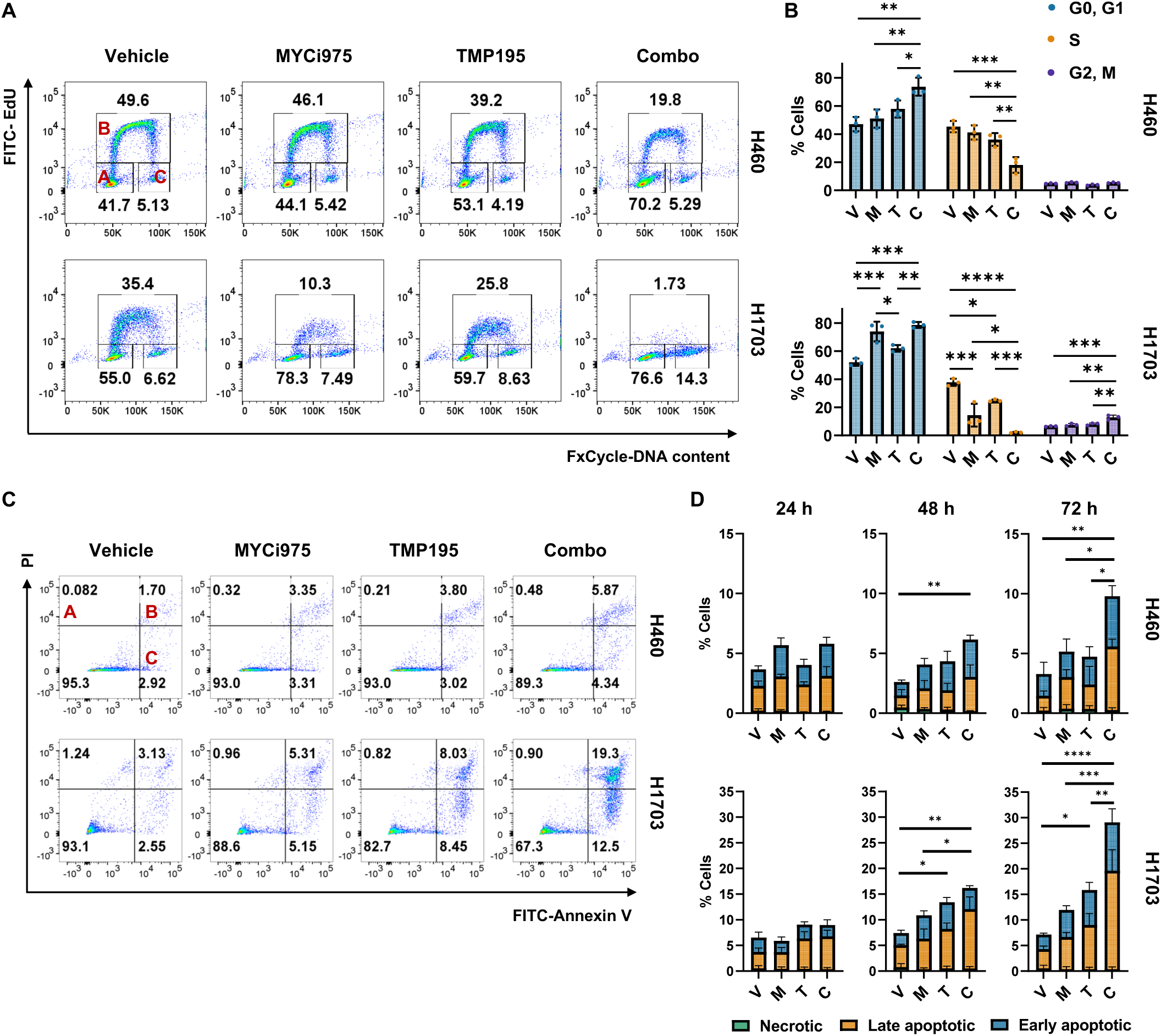
Cell cycle arrest and apoptosis induced by the combination treatment. **A** and **B,** Flow cytometry analysis of EdU and DNA staining. H460 and H1703 cells were treated with vehicle, MYCi975 (5 µM), TMP195 (25 µM), or both for 48 h, then incubated with EdU for 2 h and stained for EdU and FxCycle Violet. Representative flow cytometry dot plots illustrating the gating strategy **(A)**. A: G0/G1, B: S, C: G2/M. Percentages of cells in G0/G1, S, or G2/M phase are presented as mean ± SD **(B)**. p-values defined as: *<0.05, **< 0.01, ***<0.001, ****<0.0001; were calculated using one-way ANOVA followed by Tukey’s multiple comparisons test. n=3 biological replicates. **C,** Flow cytometry analysis of PI and Annexin V staining. H460 and H1703 cells were treated with vehicle, MYCi975 (5 µM), TMP195 (25 µM), or both for 72 h. Representative flow cytometry dot plots illustrating the gating strategy. A: Necrotic, B: Late apoptotic, C: Early apoptotic. **D,** Percentages of cells in necrotic, late apoptotic, or early apoptotic status treated with vehicle, MYCi975 (5 µM), TMP195 (25 µM), or both for 24, 48, and 72 h are presented as mean ± SD. n=3 biological replicates. p-values defined as: *<0.05, **< 0.01, ***<0.001, ****<0.0001; were calculated for necrotic + apoptotic cell percentages for each condition using one-way ANOVA followed by Tukey’s multiple comparisons test. V: vehicle, M: MYCi975, T: TMP195, C: Combo (B and D).

### MYC and cell cycle-related pathways are suppressed by combination treatment

To define transcriptional shifts facilitated by combination treatment, we performed RNA-seq on four responsive cell lines treated with vehicle, MYCi975, TMP195, or combination for 24 hours to prevent the confounder of acute cytotoxicity. Differential expression analysis on these RNA-seq data revealed markedly stronger transcriptional responses in combination drug-treated groups than in mono-treatment groups (Fig. 4, A and B). Additionally, HDAC5 was differentially upregulated in all treatment groups, whereas HDAC9 was significantly augmented only by combination treatment (Fig. 4A). Assessment of pathway-level perturbations resulting from drug treatment by Gene Set Variation Analysis (GSVA) followed by Differential Pathway Analysis revealed that the combination treatment, relative to single-agent conditions, induced a more profound suppression of cell-cycle related gene sets, such as E2F Targets, G2M Checkpoint, MYC Targets, and Mitotic Spindle (Fig. 4C). TMP195 and MYCi975 mono-treatments were also found to downregulate these pathways, but with a smaller magnitude relative to the combination treatment (Fig. 4C). GSEA on these transcriptional data demonstrated negative enrichment of cell cycle-related gene sets in combination drug-treated groups (Fig. S4A). We next examined the expression of leading-edge genes derived from GSEA on cell cycle-related pathways (Fig. S4B). This analysis showed that mono-treatments resulted in a more modest reduction in leading-edge gene expression compared to the combination treatment across all four cell lines tested (Fig. S4B). Given MYC’s role as a master regulator of cell cycle-related transcriptional networks, we assessed both MYC RNA and protein levels following drug treatment. Interestingly, while the combination treatment did not affect MYC RNA levels (Fig. 4F), MYC protein levels were significantly decreased by drug treatment in two responsive cell lines tested, H460 and H1703 (Fig. 4, D and E). However, neither mono- nor combination treatment changed MYC protein levels in two non-responsive cell lines, HCC827 and H441 (Fig. 4, D and E). Interestingly, dose-escalating to 15 µM of MYCi975 resulted in only a modest reduction of MYC protein in HCC827 and H441, while H460 and H1703 showed a complete loss of detectable MYC protein (Fig. S5A), thus suggesting an inherent resistance in these non-responding cell lines to the actions of MYCi975. To establish a causal relationship between MYC depletion and phenotypic observations, we assessed and confirmed that transient overexpression of MYC partially rescued the reduction of cell viability upon combination treatment, therefore indicating that the observed treatment effects are at least in part derived from MYC depletion (Fig. 4G and Fig. S5B).

**Figure 4.**
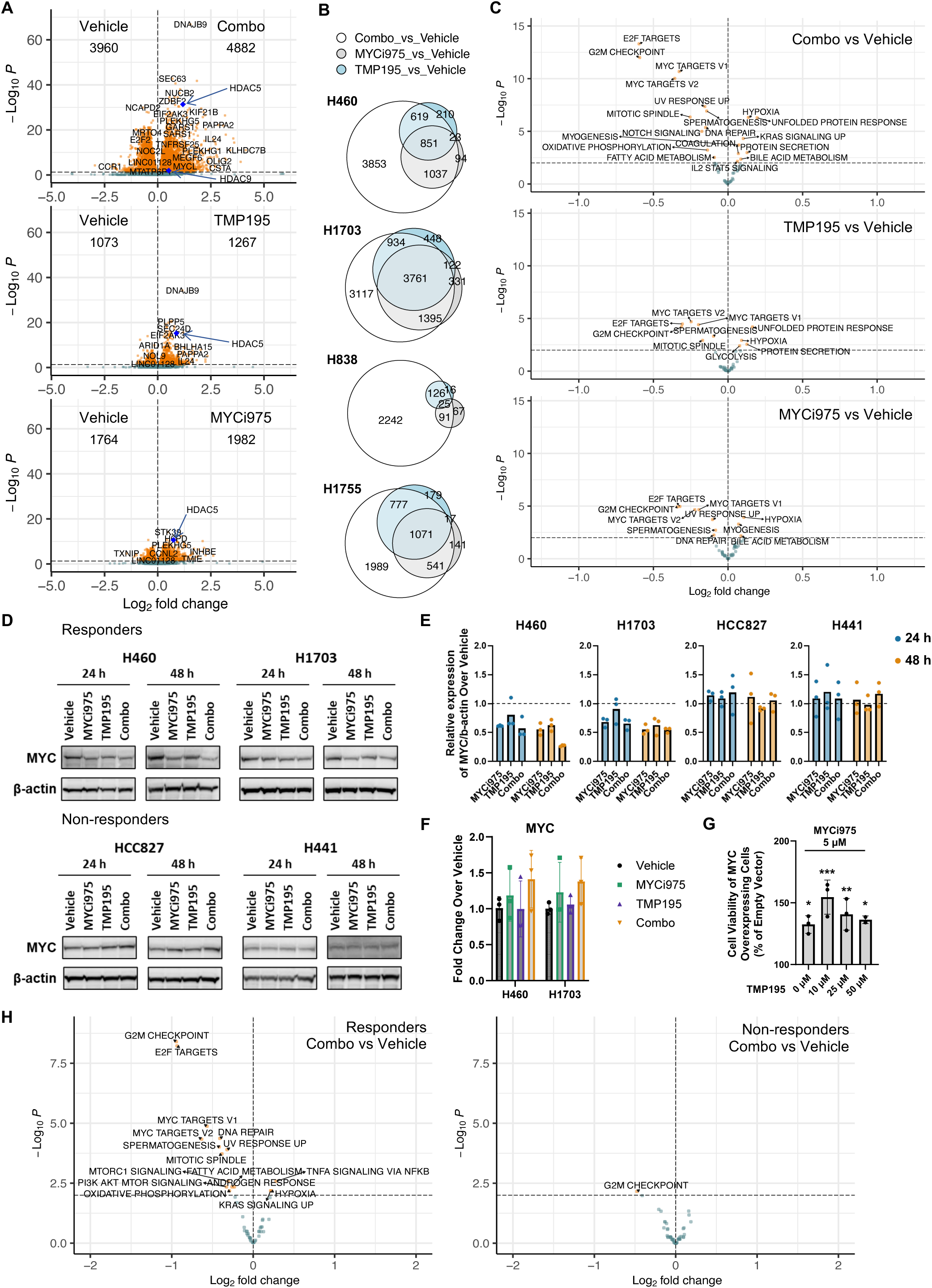
MYCi plus class IIa HDACi downregulates MYC and cell cycle-related pathways. **A,** Volcano plots for the pooled RNA-seq data (24 h treatment) from 4 responsive cell lines (H460, H1703, H838, and H1755) for Combo, TMP195, or MYCi975 vs. Vehicle. Orange dots represent differentially expressed genes with adjusted p-value < 0.05; teal dots represent genes with adjusted p-value ≥ 0.05. Blue dots represent HDAC5 or HDAC9. Each dot represents one gene. **B,** Euler plots representing the differentially expressed genes for the indicated comparisons in H460, H1703, H838, and H1755. **C,** Gene Set Variation Analysis (GSVA) with Hallmark gene sets using pooled data from 4 responsive cell lines (H460, H838, H1703, H1755) followed by differential expression analysis at the pathway level comparing Combo, MYCi975, or TMP195 vs. Vehicle. Orange dots indicate pathways with adjusted p-value < 0.01. Cells were treated with either vehicle, MYCi975, TMP195, or both for 24 h. n=4 biological replicates (A, B, and C). **D,** MYC protein levels were detected by western blotting in H460, H1703, HCC827, and H441 cells treated with indicated agents for 24 or 48 h. All immunoblots are representative of three biological replicates. **E,** Normalized MYC/b-actin ratios relative to the vehicle. n=3. **F,** RNA levels for MYC were measured by RT-qPCR in H460 and H1703 cells treated with indicated agents for 48 h. n=3 biological replicates. Mean ± SD. **G,** H460 cells were transfected with empty vector or MYC overexpression vector and treated with 5 µM MYCi975 plus indicated doses of TMP195 for 72 h. % viabilities of MYC overexpressing cells relative to the empty vector-transfected cells are depicted. Mean ± SD. p-values defined as *<0.05, **< 0.01, ***<0.001 were calculated using a t-test for indicated condition vs. empty vector. n=2 or 3. **H,** Gene Set Variation Analysis (GSVA) with Hallmark gene sets using pooled data from 2 responsive cell lines (H460, H1703) or 2 non-responsive cell lines (HCC827, H441) followed by differential expression analysis at the pathway level comparing Combo vs. Vehicle. Orange dots indicate pathways with adjusted p-value < 0.01. Cells were treated with either vehicle or MYCi975 + TMP195 for 48 h. n=2 biological replicates.

Based on the robust reduction of MYC protein levels induced by combination treatment in responders (H460 and H1703) at 48 hours post-treatment initiation but not in two non-responders (HCC827 and H441), we sought to investigate transcriptional alterations in these four cell lines at this time point. These data revealed that combination treatment induced more substantial transcriptional shifts in responders than non-responders, as evidenced by the absolute number of DEGs (Fig. S5C). Pathway analysis by GSVA revealed that responsive cell lines presented robust suppression of cell cycle-related pathways, whereas changes in those pathways were more muted in non-responders (Fig. 4H). RT-qPCR validated combination treatment-induced repression of critical genes involved in cell cycle progression, including AURKA, AURKB, PLK1, PLK4, CDC25A, and E2F2 upon combination treatment for 48 hours (Fig. S5D). Consistently, this suppression of cell cycle-related gene expression was more potent in responsive cell lines (H460 and H1703) compared to non-responders (HCC827 and H441), suggesting that transcriptional shifts may define treatment effects (Fig. S5D). These data collectively demonstrate that combination treatment causes suppression of cell cycle progression-related pathways with an associative depletion of MYC protein, which has a causal relationship with the observed phenotype.

Considering that MYC has been reported to alter chromatin structure, changes in chromatin accessibility upon the combination treatment in H460 and H838 cells were assayed through ATAC-sequencing.(67, 68) Interestingly, in both cell lines, more significant numbers of differentially accessible regions (DARs) were driven by combination drug treatment relative to the mono-treatments (Fig. 5, A and B and Fig. S6, A to C). Genomic Regions Enrichment of Annotations Tool (GREAT)(69) revealed that downregulated DARs were enriched for cell cycle-related pathways in both H460 and H838 cell lines upon combination treatment (Fig. 5C and Fig. S6D). We integrated RNA-seq and ATAC-seq data by identifying activated or suppressed transcription factors (TF) in both datasets to elucidate chromatin changes associated with transcriptional alterations. We first assessed positively or negatively enriched TF motifs in combination-treated cells compared to the vehicle group using HOMER with ATAC-seq data. Next, GSEA on TF target gene sets (C3:TFT) with RNA-seq data identified significantly upregulated or downregulated TFs upon combination treatment. These analyses revealed that MYCi975 plus TMP195 activated TFs involved in stress response and inflammation, such as ATF3 and AP-1, while E2F TFs were repressed upon combination treatment (Fig. 5, D and E and Fig. S6, E and F).

**Figure 5.**
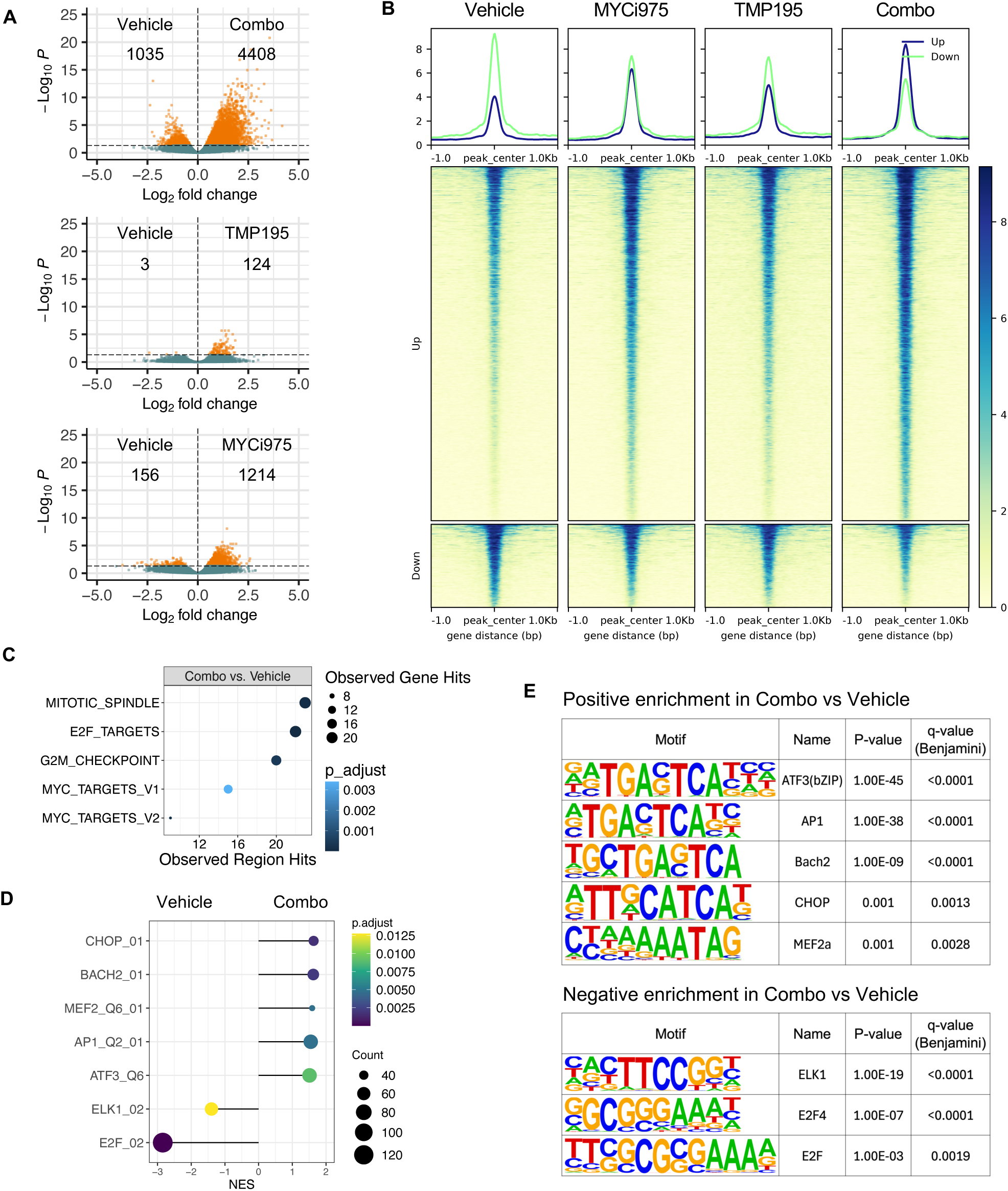
Combination treatment induces large-scale changes in chromatin accessibility. **A,** Volcano plots of ATAC-seq consensus peaks. Orange dots represent differentially accessible regions with adjusted p-value < 0.05. Each dot represents one peak. H460 cells were treated with vehicle, MYCi975 (5 µM), TMP195 (25 µM), or both for 24 h. n=4 biological replicates. **B,** Heatmaps representing ATAC-seq peaks within differentially accessible regions (Combo vs. Vehicle) for indicated samples in H460 cells. **C,** GREAT (the Genomic Regions Enrichment of Annotations Tool) analysis on cell-cycle related pathways in H460 cells. **D,** GSEA with C3:TFT pathways based on differential expression analysis of combo vs. vehicle-treated H460 samples. **E,** Lists of transcription factors (TF) with motifs that are positively or negatively enriched in the combo group compared to the vehicle group in ATAC-seq data and with their target gene sets enriched in RNA-seq in H460.

### Mitochondrial dysfunction is induced by combination of MYC and class IIa HDAC inhibition

MYC is a crucial regulator of mitochondrial biology, including oxidative phosphorylation, mitochondrial biogenesis, mtDNA replication, and transcription.(70–73) Thus, we hypothesized that mitochondrial dysfunction may result from combination treatment. Pathway analysis by GSVA on mitochondrial gene signatures demonstrated that combination treatment suppressed multiple mitochondrial transcriptional programs, including mitochondrial DNA replication and maintenance, TCA cycle, and OXPHOS subunits (Fig. 6A). In contrast, mono-treatments induced only modest perturbations on mitochondrial pathways (Fig. 6A). Next, we compared changes in mitochondrial pathways in responders and non-responders at 48 hours post-combination treatment. Interestingly, no mitochondrial pathways were significantly altered in non-responders (Fig. 6B). In contrast, pathways involved in critical mitochondrial functions, including mtDNA and OXPHOS, were downregulated in responding cell lines (Fig. 6B). Based on these data demonstrating transcriptional suppression of mitochondrial pathways, we next examined mitochondrial membrane potential, which is a measure of overall mitochondrial activity.(74) Interestingly, we observed a loss of mitochondrial membrane potential upon MYCi plus class IIa HDACi in both H460 and H1703 cells (Fig. 6, C and D). Particularly in H1703, a cell line that showed strong apoptosis induction by the combination treatment (Fig. 3), we observed mitochondrial membrane potential reduction at 24 hours post-treatment initiation, which preceded the apoptotic response observed at 48 hours. Disrupted mitochondrial membrane potential was accompanied by an augmentation of mitochondrial localized superoxide driven by MYCi975 treatment (Fig. 6, E and F). In contrast, no significant change was detected in cytosolic superoxide during the drug treatment (Fig. 6, E and F). Finally, we assessed whether ROS contributes to treatment effects derived from the combination treatment using N-acetylcysteine (NAC), a ROS scavenger. These data revealed that treatment with NAC rescued the combination therapy-associated viability defect in a dose-dependent manner (Fig. 6G). Collectively, these findings show that combination treatment induces mitochondrial dysfunction and associated mitochondrial ROS induction, which are at least partially responsible for combination treatment effects.

**Figure 6.**
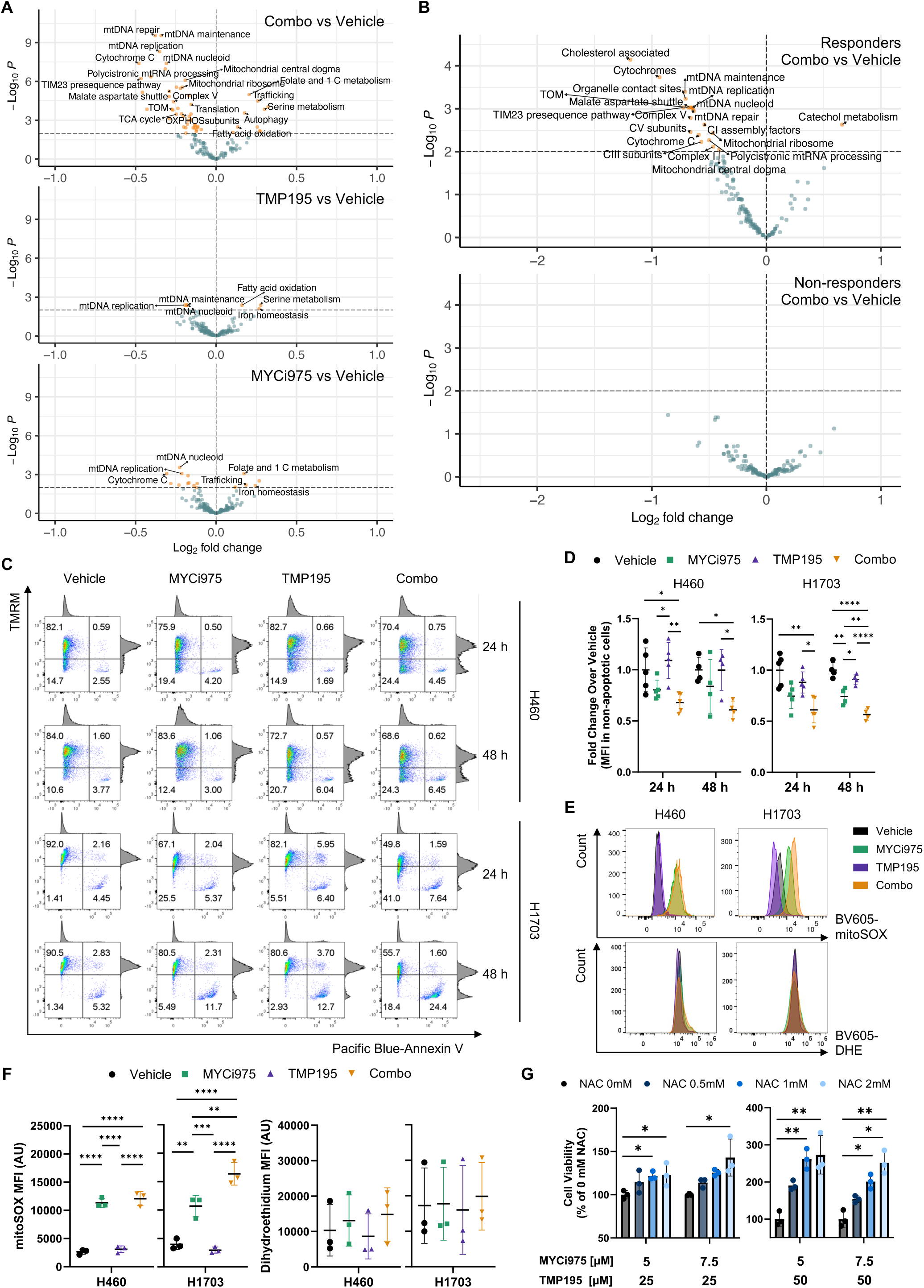
MYCi plus class IIa HDACi causes mitochondrial dysfunction and mitochondrial ROS induction. **A,** GSVA with mitoCarta 3.0 gene sets using pooled data from 4 responsive cell lines (H460, H838, H1703, H1755) followed by differential expression analysis at the pathway level comparing Combo, MYCi975, or TMP195 vs. Vehicle. Orange dots indicate pathways with adjusted p-value < 0.01. Cells were treated with either vehicle, MYCi975, TMP195, or both for 24 h. n=4. **B,** GSVA with mitoCarta 3.0 gene sets using pooled data from 2 responsive cell lines (H460, H1703) or 2 non-responsive cell lines (HCC827, H441) followed by differential expression analysis at the pathway level comparing Combo vs. Vehicle. Orange dots indicate pathways with adjusted p-value < 0.01. Cells were treated with either vehicle or MYCi975 + TMP195 for 48 h. n=2. **C** and **D,** Flow cytometry analysis of Tetramethylrhodamine (TMRM), a mitochondrial membrane potential indicator, and Annexin V. H460 and H1703 cells were treated with indicated agents for 24 or 48 h. Representative flow cytometry dot plots illustrating the gating strategy **(C)**. Fold change of TMRM mean fluorescence intensity (MFI) in Annexin V-negative cells over vehicle **(D)**. n=5 for 24 h, n=4 for 48 h. **E** and **F,** Flow cytometry analysis of mitoSOX, a mitochondrial superoxide indicator, and DHE (Dihydroethidium), a cytosolic superoxide indicator. H460 and H1703 cells were treated with indicated agents for 24 h. Histogram for mitoSOX or DHE signals **(E)** and MFI **(F)**. n=3 biological replicates. **G,** H460 cells were treated with NAC and indicated drugs for 72 h. % viabilities of NAC-treated cells relative to untreated cells are depicted. p-values for indicated condition vs. 0 mM NAC are depicted. n=3 biological replicates. Data are presented as mean ± SD (D, F, and G). p-values defined as *<0.05, **< 0.01, ***<0.001, ****<0.0001; were calculated using one-way ANOVA followed by Tukey’s multiple comparisons test (D, F, and G).

### Combination of MYC and class IIa HDAC inhibition reduces tumor burden in vivo

To assess the cancer cell-intrinsic effects of the combination treatment, we first utilized an H460 xenograft mouse model. In this model, we observed a significant reduction in tumor volume in mice treated with the combination of MYCi975 and TMP195 compared to vehicle-treated mice (Fig. S7, A and B). In contrast, mono-treatments failed to demonstrate a statistically significant reduction in tumor growth relative to the vehicle controls (Fig. S7, A and B). To enhance the translational applicability of our study, we evaluated therapeutic efficacy in a patient-derived xenograft (PDX) model harboring a KRAS G12D mutation. Similar to the pattern observed in the H460 xenograft model, combination treatment resulted in a significant reduction of tumor volume in the PDX model, which was most profound in combination therapy-treated mice (Fig. S7, C and D). Finally, we assessed the therapeutic efficacy in the murine syngeneic adenocarcinoma Lewis Lung Carcinoma (LLC1) model, which was chosen to evaluate the perturbation of immune populations and signaling pathways driven by treatment in an immune-competent and clinically relevant setting.(75) In this model, tumor growth was significantly reduced by combination treatment compared to the vehicle, MYCi975, and TMP195 groups (Fig. 7, A and B). Importantly, no significant weight loss was noted in any models tested (Fig. S7, E to G).

**Figure 7.**
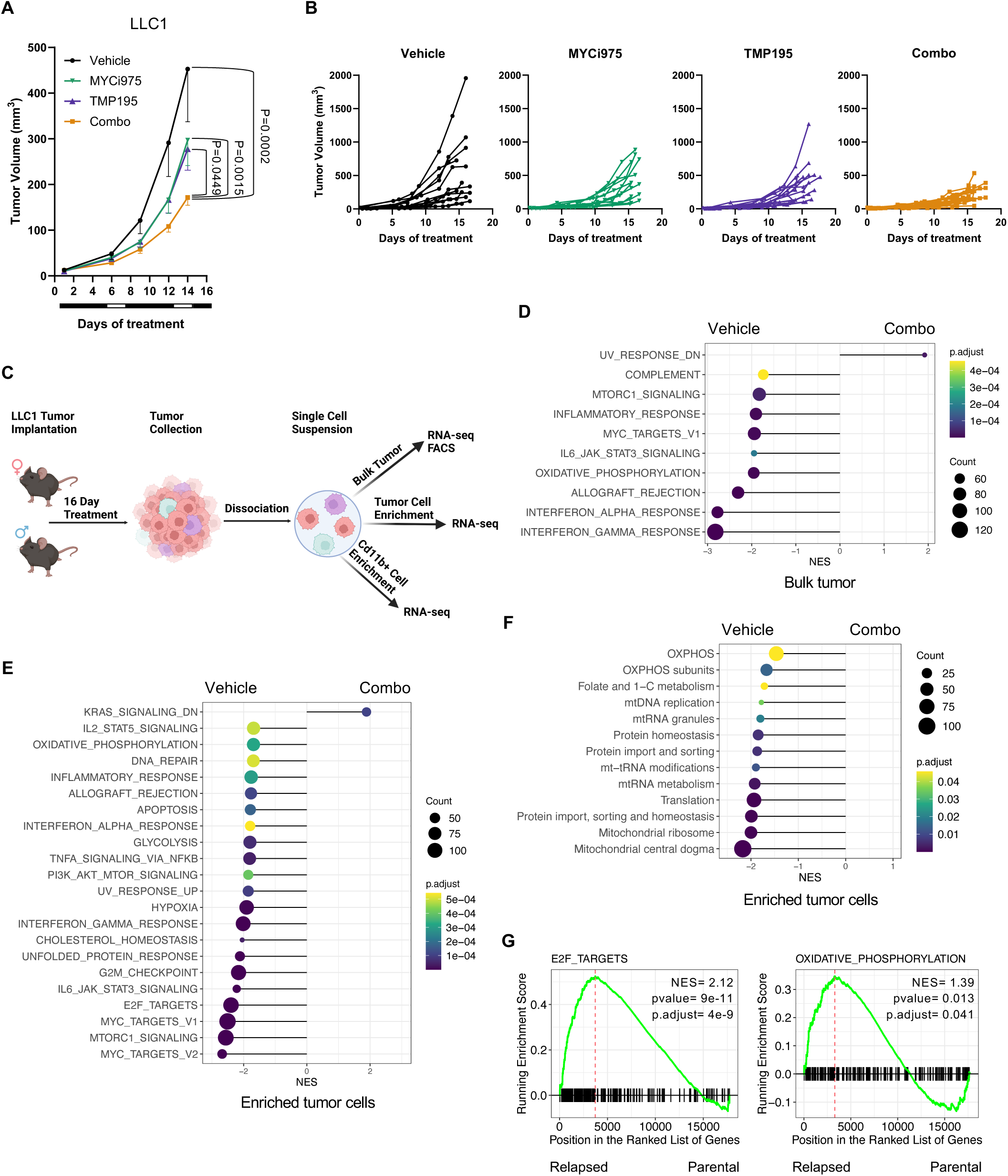
MYCi plus class IIa HDACi reduces tumor growth in vivo. **A** and **B,** Mean volumes of LLC1 tumors implanted in C56BL/6 mice treated with the drugs as indicated **(A)**. Spider plots depict individual tumor growth **(B)**. n=12 for vehicle, MYCi975, TMP195, n=15 for Combo. Treatment schema below each plot: black indicates on-treatment, and white indicates off-treatment. Mean ± SEM. Linear mixed model by the Restricted maximum likelihood (REML) approach was used to derive p-values. **C,** LLC1 tumor model molecular correlate schema. **D,** GSEA with Hallmark pathways based on differential expression analysis with bulk tumor from combo vs. vehicle-treated mice. n=4 for vehicle and n=3 for combo. **E** and **F,** GSEA with Hallmark **(E)** or MitoCarta 3.0 **(F)** pathways based on differential expression analysis with enriched tumor cells from combo vs. vehicle-treated mice. n=4. LLC1 tumor-bearing mice were treated with either vehicle or combo for 16 d (D, E, and F). **G,** GSEA with E2F Targets and Oxidative Phosphorylation Hallmark pathways based on differential expression analysis with Cd45^−^ tumor cells derived from ICB (Immune Checkpoint Blockade)-relapsed vs. parental LLC (GSE246922). Running enrichment scores and the distribution of gene sets in the ranked list of genes are depicted.

Considering the robust transcriptional effects underpinning in vitro drug responses, we pursued bulk-tumor-, tumor-cell-enriched-, and Cd11b-enriched-RNA-seq on drug-treated LLC1 tumors to further explore molecular correlates associated with the observed in-vivo phenotype (Fig. 7C). Assessment of bulk-tumor transcriptomic data revealed a limited number of DEGs across all conditions assessed (Fig. S8A). Pathway analysis of bulk-tumor-RNA-seq data defined a consistent decrease in oxidative phosphorylation across all treatment groups (Fig. 7D and Fig. S8B). In contrast, immune signaling pathways, such as interferon alpha response, interferon gamma response, and IL6/JAK/STAT3 signaling, were only downregulated in combination treatment conditions, while mono-treatment activated these pathways (Fig. 7D and Fig. S8B). Assessment of tumor infiltrative populations by fluorescence-activated cell sorting (FACS) failed to define any high-level changes in tumor cellularity resulting from treatment (Fig. S9, A and B). Evaluation of tumor-cell-enriched-RNA-seq defined a modest set of DEGs of nearly equal magnitude across all treatment conditions (Fig. S9C). Pathway analysis of these tumor cell-enriched transcriptional data revealed a profound decrease in MYC targets and cell cycle-associated pathways in all treatment groups (Fig. 7E and Fig. S9D). Assessment of mitochondrial pathway perturbations using MitoCarta 3.0 gene sets uncovered a conserved pattern of negative enrichment across oxidative phosphorylation, mitochondrial metabolism, and central dogma-associated pathways in TMP195 and combination-treated enriched tumor cells, whereas MYCi975 treatment resulted in no significant enrichment of these pathways (Fig. 7F and Fig. S9E).

To further contextualize the significance of our LLC1 tumor RNA-seq findings, we reanalyzed RNA-seq data from Cd45^−^ populations in anti-PD-1 and anti-CTLA-4 resistant LLC1 to discern whether any of the noted pathways were operative in the acquisition of treatment resistance (GSE246922) (Fig. 7G and Fig. S9, F and G).(76) Analysis of these data revealed a significant increase in oxidative phosphorylation and cell cycle pathways in treatment-resistant disease (Fig. 7G and Fig. S9F). Additionally, several mitochondria-specific pathways were also augmented in relapsed disease, although none were determined to be statistically significant (Fig. S9G).

As tumor immune microenvironment is cellularly heterogeneous, we sought to determine the cellularity of Cd45^+^ cell populations in LLC1 tumors by querying publicly deposited single-cell RNA-seq data (GSE169688).(77) Evaluation of these data revealed Cd11b^+^ cells to be the dominant immune cell resident population in LLC1 tumors (ref.(77) and Fig. S10A); we, therefore, selected this population for further molecular characterization. Interrogation of Cd11b-enriched-RNA-seq uncovered a more profound transcriptional response from therapy relative to bulk or tumor cell-enriched populations (Fig. S10B). Interestingly, TMP195 and combo groups showed demonstrable repression of inflammatory pathways, while MYC inhibition alone augmented these pathways (Fig. S10C). Critically, cell cycle and oxidative phosphorylation-associated pathways were not significantly impacted in Cd11b^+^ cells by combination treatment, thus suggesting a tumor cell-specific effect (Fig. S10C). Given the known role of Cd11b^+^ cells in potentiating neovascularization,(78–81) we sought to define whether perturbation of vascularization processes was evident at the pathway level. Targeted assessment of vascularization and endothelial cell-associated pathways in tumor-infiltrating Cd11b^+^ cells revealed significant negative enrichment of pathways involved in vasculature development and establishment of endothelial barrier, most apparent in combination-treated tumors (Fig. S10, D and E). Furthermore, cell type signature gene sets and marker genes associated with endothelial cells were transcriptionally repressed in response to therapy, which was most profound for combination-treated bulk tumor samples (Fig. S10, F and G).

## Discussion

Herein, we define a therapeutic approach combining MYC and class IIa HDAC inhibitors for NSCLC treatment. Notably, response to the combination treatment was associated with the downregulation of MYC target and cell cycle-related gene expression and post-transcriptional MYC repression. In addition, MYCi plus class IIa HDACi causes suppression of mitochondrial pathways and activity and augments mitochondria-localized ROS. Mechanistically, MYC depletion and oxidative stress were found to drive combination drug efficacy.

MYC targeting has been intensely studied as a cancer treatment strategy based on the frequent dysregulation of MYC signaling in cancer and promising results from preclinical research investigating MYC suppression in mouse cancer models.(26, 40) However, no direct MYC targeting agents are currently approved for cancer treatment due to difficulties in pharmacological inhibition of MYC.(40) Nevertheless, diverse approaches to inhibit MYC have been tested in preclinical and clinical settings. One such strategy involves MYC suppression by silencing its expression. For example, DCR-MYC, a siRNA targeting MYC, has been tested in a phase I clinical trial (NCT02110563) and was found to be well-tolerated, although no further studies have been conducted using this drug.(82) Additionally, G-quadruplex stabilizing agents, such as APTO-253 and CX-5461, which are known to suppress MYC transcription, have been evaluated in phase I clinical trials.(83–85) Although APTO-253 was generally well-tolerated and shown to reduce MYC expression levels, its clinical development was discontinued per the company’s decision.(84) CX-5461, which was also well-tolerated, demonstrated clinical responses in 14% of patients with germline DNA repair abnormalities.(83) Moreover, MYC can be targeted by promoting its proteasomal degradation through PLK1 or AURKA inhibitors, both of which have been tested in several clinical trials.(86, 87) However, PLK1 or AURKA-targeting agents have demonstrated limited anti-cancer efficacy as mono-therapies in the clinic. These indirect MYC inhibiting approaches have limitations since multiple layers of context-dependent mechanisms at the transcriptional, translational, mRNA, and protein levels control MYC activity. In addition, due to the prominent role of MYC in cancer biology, resistance to these agents might result from MYC re-expression via various compensatory mechanisms. For example, tumors treated with MYC silencing drugs can acquire mutations in MYC, enhancing the protein stability and leading to drug resistance. Given these drawbacks, drugs that perturb multiple aspects of MYC biology simultaneously, such as MYCi975, might prove more efficacious. Moreover, considering the significance of MYC in cancer progression, combination treatment strategies might be needed to achieve durable responses and target pro-survival compensatory mechanisms initiated post-MYC inhibition.(88, 89) Here, we propose class IIa HDAC inhibitors as a combination partner for MYC targeting agents to derive prolonged and robust treatment efficacy in NSCLC.

Given the high heterogeneity of drug response in cancer, it is critical to identify molecular markers linked to drug sensitivity. This study defined transcriptional, genomic, and proteomic profiles correlated with response, which may help identify patient populations primed to respond to our drug regimen. Specifically, we elucidated that differential responses to the combination treatment are associated with altered MYC targets, oxidative phosphorylation, ROS, and mitochondrial pathways transcriptionally. Furthermore, EGFR-altered cancer cell lines were non-responsive to MYCi plus class IIa HDACi, while cell lines with genomic alterations in KRAS/STK11 genes were more likely to be sensitive to the combination treatment. MYC protein levels correlate with combination treatment responses, consistent with previous studies reporting the association between MYC levels and treatment response to MYC inhibition.(38, 90, 91) Based on these data, genomic profiling of clinical tumor samples paired with immunohistochemistry for MYC could be beneficial for patient selection. These findings on molecular vulnerability shed light on potential mechanisms of combination treatment and correlative biomarkers that predict response to our treatment paradigm.

Aberrant regulation of mitochondrial function is a well-established hallmark of cancer inherently involved in tumorigenesis. Mitochondria supply adenosine triphosphate (ATP), macromolecules, and metabolite precursors, thus allowing cancer cells to meet their high metabolic demands.(92, 93) Mitochondria are significant sources of ROS and redox molecules in multiple antioxidant pathways controlling redox homeostasis.(94) ROS levels are known to be elevated in cancer cells, which in turn activate oncogenic pathways.(95) These augmented ROS levels, however, might provide a therapeutic opportunity to raise oxidative stress to a degree where survival is untenable. Importantly, it has been proposed that cancer cells are more susceptible to mitochondrial perturbation than normal cells due to metabolic reprogramming, thereby presenting a therapeutic window for mitochondrial targeting.(96, 97) Despite intense research on mitochondrial inhibition as a cancer treatment modality, clinical benefits from existing approaches have remained elusive.(92) One potential method for disrupting mitochondrial function is through MYC suppression. MYC promotes respiration and macromolecule synthesis in cancer by enhancing mitochondrial biogenesis and metabolism to fuel rapid tumor growth.(93) Nevertheless, how MYC suppression in cancer affects mitochondria and how this mitochondrial dysregulation contributes to anti-tumor effects remain unclear. This study found that MYCi plus class IIa HDACi suppresses mitochondrial transcriptional pathways and activity. Notably, apoptosis is preceded by a loss of mitochondrial membrane potential and an augmentation of mitochondrial ROS, which suggests a link between mitochondrial perturbation and therapeutic effects. Moreover, cancer cells with higher ROS and mitochondrial pathways, such as OXPHOS, mitochondrial central dogma, and metabolism, were more sensitive to the combination treatment, implicating mitochondrial phenotypes in treatment effects. Thus, this work connects MYCi plus class IIa HDACi-induced mitochondrial dysfunction and cell-intrinsic anti-cancer effects in NSCLC.

Our present in vivo studies demonstrate the anti-tumor efficacy of MYCi plus class IIa HDACi in xenograft and syngeneic mouse cancer models, illustrating the translational potential of this drug paradigm for NSCLC treatment. Importantly, we observed marked anti-tumor effects from the combination treatment in the syngeneic LLC1 model. This model is intrinsically resistant to immune checkpoint inhibitors, including anti-CTLA4 and anti-PD-1.(98) Therefore, our data suggests that MYCi plus class IIa HDACi might be one approach to target NSCLC unresponsive to current immunotherapies.(99) Furthermore, augmented OXPHOS in tumors has been implicated in resistance to both chemotherapy and immune checkpoint blockade therapy.(71, 100–102) Notably, a recent study has reported post-treatment upregulation of the OXPHOS pathway in NSCLC patients with acquired resistance to immune blockade therapy,(76) thus highlighting the potential therapeutic benefit derived from OXPHOS suppression by our combination drug approach.

Additionally, we have shown the suppression of endothelial cell signatures and associated downregulation of pathways involved in vasculature/endothelium development in tumor-infiltrating Cd11b^+^ populations. Of note, a class IIa HDAC inhibitor used in this study, TMP195, has been demonstrated to improve the integrity of tumor vasculature in a breast cancer model.(57) Given that myeloid cells, such as tumor-associated macrophages and myeloid-derived suppressive cells, play essential roles in tumor angiogenesis,(78–81) our combination paradigm could be one way to target this process. Although the combination treatment did not significantly alter the composition of immune populations in LLC1 tumors, further study is warranted to determine whether there are changes in the functionalities of immune populations and how these could contribute to treatment efficacy.

In summary, we propose a combination drug paradigm, which exerts robust anti-tumor effects in NSCLC with basally high MYC and mitochondrial activity. Both MYC depletion and oxidative stress from mitochondria-localized ROS partially drive these treatment effects. Furthermore, this combination treatment reduces tumor burden across multiple lung cancer murine models, indicating the translational potential of our study for the treatment of NSCLC.

## Methods

### Non-Small Cell Lung Cancer Cell Lines and Cell Culture

Cell lines used in the study were acquired from ATCC. Human NSCLC cell lines were cultured in RPMI1640 media (Corning) containing 10% v/v FBS (Gemini Bio) at 37°C and 5% CO_2_. Human NSCLC cell lines: NCI-H1755, NCI-H520, NCI-H1650, NCI-H441, NCI-H661, NCI-H596, HCC4006, NCI-H1703, NCI-H838, NCI-H2170, NCI-H1792, NCI-H460, NCI-H23, A549, NCI-H1299, HCC827, NCI-H1869, NCI-H226. Lewis Lung Carcinoma (LLC1) cells were cultured in DMEM media (Corning) containing 10% v/v FBS (Gemini Bio) at 37°C, 5% CO_2_.

### Drug Treatment in vitro and in vivo

For MYCi975 plus TMP269 in vitro treatment, MYCi975 (Selleck Chemicals) and TMP269 (ApexBio) were dissolved in DMSO to concentrations of 1 mM, 5 mM, 10 mM, 25 mM, and 50 mM and stored at −20°C. For MYCi975 plus TMP195 in vitro treatment, MYCi975 (DC Chemicals) and TMP195 (DC Chemicals) were dissolved in DMSO to concentrations of 1 mM, 5 mM, 10 mM, 25 mM, and 50 mM and stored at −20°C. N-acetylcysteine (NAC) was dissolved in water at 200 mM prior to each treatment. NAC was pretreated for 1 h for the rescue experiments, followed by MYCi975 and TMP195 treatment. For in vivo treatment, MYCi975 (DC Chemicals) and TMP195 (DC Chemicals) were freshly prepared in 10% DMSO + 30% PEG300 + 5% Tween 80 + 55% PBS.

### CellTiter-Glo Luminescent Cell Viability Assay and Viability Data Analysis

In white-walled 96 well plates, cell lines were plated at the following densities: NCI-H1755 (2.0×10^3^), NCI-H520 (2.0×10^3^), NCI-H1650 (2.0×10^3^), NCI-H441 (4.0×10^3^), NCI-H661 (2.0×10^3^), NCI-H596 (2.0×10^3^), HCC4006 (2.0×10^3^), NCI-H1703 (2.0×10^3^), NCI-H838 (1.0×10^3^), NCI-H2170 (2.0×10^3^), NCI-H1792 (2.0×10^3^), NCI-H460 (1.0×10^3^), NCI-H23 (2.0×10^3^), A549 (1.0×10^3^), NCI-H1299 (1.0×10^3^), HCC827 (4.0×10^3^), NCI-H1869 (4.0×10^3^), NCI-H226 (2.0×10^3^). At 1 day post-plating, cells were treated with DMSO or combinations of MYCi975 plus TMP269/TMP195 for 72 hours and subjected to CellTiter-Glo Luminescent Cell Viability Assay (Promega, G7572). Cell lysis was induced by adding CellTiter-Glo reagent and shaking the plates for a minute. Plates were incubated at room temperature for 10 minutes, and luminescence was measured using the POLARStar OPTIMA (BMG LABTECH). GraphPad Prism was used to calculate Area Under the Curve (AUC) values from the MYCi975 dose-response curves. For normalized proliferation graphs, the luminescence data collected from two or three independent experiments were normalized, and drug doses were log-transformed. Using GraphPad Prism, these normalized, log-transformed data were analyzed by 4-parameter nonlinear regression to generate normalized MYCi975 log dose-response curves. Combenefit software was used for the Bliss model-based synergy calculation.

### Class IIa HDAC Glo Assay

Cell lines were plated in white-walled 96 well plates at a concentration of 8.0×10^3^. At 1 day post-plating, cells were treated with indicated drugs for 24 hours, and HDAC-Glo™ Class IIa Assay (Promega) was performed according to the manufacturer’s protocol. The luminescence was measured using the POLARStar OPTIMA (BMG LABTECH).

### CCLE Genomic Alteration Analysis and Oncoprint

CCLE Mutation data for the cell lines were acquired from the DepMap portal. Significant mutated genes in either LUAD or LUSC reported in Campbell, J. D., et al. were gathered to annotate hotspot mutations.(17) Among these genes, those with non-silent and hotspot (TCGA or COSMIC) mutations in at least two cell lines were selected for the Oncoprint plot. Genes with significantly recurrent copy number losses in LUAD or LUSC were gathered to annotate deletion. Among these genes, those with an absolute copy number of 0 in at least two cell lines were depicted in the Oncoprint plot. Genes with significantly recurrent copy number gains in LUAD or LUSC were gathered to annotate amplification. Among these genes, those with an absolute copy number > 5 and a copy number ratio > 2 in at least two cell lines were depicted.

### CCLE Proteomics Analysis

CCLE proteomics data for the 14 cell lines (9 responders, 5 non-responders) was available on the DepMap portal. Normalized protein levels for MYC, HDAC4, HDAC5, HDAC7, and HDAC9 were gathered. Cell lines with a missing value for each protein were not included in the analysis or visualization, as suggested by Nusinow, D. P., and S. P. Gygi. (2020).(bioRxiv 2020.02.03.932384)

### Cell Cycle Assay and Apoptosis Assay using Flow Cytometry

Cell cycle assay was performed using Click-iT EdU Alexa Fluor 488 Flow Cytometry Assay Kit (Invitrogen, C10425) according to the manufacturer’s protocol. In brief, cells were treated with 10 µM EdU for 2 hours. After fixation and permeabilization, cells were subjected to Click-iT reaction and DNA content staining with FxCycle Violet. The fluorescent signals generated by Click-iT EdU labeling, and DNA staining were detected by flow cytometry using BD LSRFortessa. Apoptosis assay was performed using the Dead Cell Apoptosis Kits with Annexin V for Flow Cytometry (Invitrogen, V13241).

### RNA-seq Library Generation for NSCLC Cell Lines

H460, H1703, H838, and H1755 cells were treated with vehicle, 5 µM MYCi975, 25 µM TMP195, or both for 24 hours or 48 hours, and total RNA was isolated using the NucleoSpin RNA purification mini kit (Macherey-Nagel, 740955.50). Biological replicates were prepared from independent experiments. NanoDrop ND-1000 and Bioanalyzer (Agilent Technologies) were used to measure RNA concentration and integrity, respectively. Following the manufacturer’s protocol, RNA-seq libraries were prepared using the SMARTer Stranded Total RNA Sample Prep Kit (Takara, 634875). Bioanalyzer and NEBNext Library Quant Kit for Illumina (NEB, E7630) were used to assess the quality and quantity of each library, respectively. Prepared libraries were sequenced on the Illumina NOVAseq 6000 instrument for paired-end reads.

### qRT-PCR

1ug RNA from drug-treated cell lines was converted into cDNA using qScript cDNA Supermix (Quantabio, 95048). qRT-PCR was performed on cDNA samples using SsoAdvanced Universal SYBR Green Supermix (Bio-Rad, 1725275). TBP and UBC were used as reference genes for normalization. Primer sequences are shown in Supplementary Table 2.

### Western Blotting

Cells were lysed in RIPA buffer (Sigma, R0278) with protease and phosphatase inhibitors (Thermo Scientific, 78440). Protein concentrations were measured by Bradford Assay (Thermo Fisher, 23246). Equal protein quantities were subjected to SDS-PAGE with 4%–12% Bis-Tris BOLT gel (Life Technologies) and transferred to PVDF membrane (Millipore). Membranes were blocked in 10% milk/TBST and immunoblotted with the following antibodies: rabbit monoclonal c-MYC (Cell Signaling, 1:1000) and Mouse monoclonal anti-b-Actin (Sigma-Aldrich, 1:5000). The loading control, anti-b-Actin antibodies were applied after membrane stripping with stripping buffer (Thermo Scientific, 46430). Signal intensities of immunoblots were analyzed using ImageJ.

### Cell Viability Assay with MYC Overexpressing H460 Cells

pcDNA3-cMYC (Addgene, #16011) was reverse-transfected in H460 cells using Lipofectamine 3000 Transfection Reagent (Invitrogen, # L3000001) following the manufacturer’s protocol. Briefly, H460 cells were plated and concurrently transfected with pcDNA3-cMYC. On day 1, cells were replated in a white-walled 96 well, and on day 2, cells were treated with 200 µg/ml zeocin and combination drugs. On day 5, the cell titer glo assay was performed as described.

### ATAC-seq Library Generation and ATAC-seq Data Analysis

The Omni-ATAC method (Corces, M. Ryan, et al. 2017) was used to prepare the ATAC-seq libraries.(103) Experiments for four biological replicates were performed independently. In brief, H460 and H838 were treated with vehicle, 5 µM MYCi975, 25 µM TMP195, or both for 24 hours. 50,000 viable cells from each sample were collected and lysed with cold lysis buffer containing 0.1 % v/v NP-40, 0.1 % v/v Tween-20, and 0.01 % v/v Digitonin. Following the centrifuging of the cell lysate, the pellet (nuclei) was incubated with Tn5 Transposase (Illumina Tagment DNA TDE1 Enzyme and Buffer Kits, 20034197) for 30 minutes at 37°C on a thermomixer at 1,000 rpm. DNA was extracted from the lysate, followed by PCR amplification with the number of cycles determined by qPCR. ATAC-seq libraries were purified with AMPure XP beads (Beckman Coulter, A63880), and the quantity and quality of each library were measured. These libraries were sequenced on the Illumina NOVAseq 6000 instrument for paired-end reads. Raw FASTQ files were trimmed using Trimmomatic v0.39 and aligned using Bowtie2 v2.3.5.1 to the GENCODE GRCh38 genome. In the resulting BAM files, reads that are unmapped, mate-unmapped, not primary alignment, failing platform, PCR duplicates, mapping to chrM, or multi-mappers (MAPQ <20) were removed. MACS3 was used for peak-calling, and Diffbind(104) was used for obtaining consensus peak sets, counting reads for the peaks, normalization, and Differentially Accessible Region (DAR) analysis (EdgeR). DeepTools was used to generate heatmaps representing ATAC-seq peaks within differentially accessible regions (Combo vs. Vehicle) for a representative replicate. GREAT analysis was performed to identify the functional enrichment of DARs. For motif analysis, peaks from 4 replicates were merged, and HOMER v4.11 was used to identify enriched motifs in genomic regions (findMotifsGenome.pl).

### Mitochondrial Membrane Potential and Apoptosis Assay using Flow Cytometry

Following the manufacturer’s protocol, the mitochondrial membrane potential was assessed using the MitoProbe TMRM Assay Kit for Flow Cytometry (Thermo Scientific, M20036). In brief, cells were prepared in complete medium at 1.0×10^6^ cells/ml and incubated with 20 nM TMRM reagent at 37°C for 30 minutes in the presence of 5% CO_2_. Cells were washed with PBS and resuspended in Annexin Binding Buffer (Invitrogen, V13246). Apoptotic cells were stained with Annexin V Pacific Blue conjugate (Invitrogen, R37177). Fluorescent signals generated by TMRM and Annexin V were detected by flow cytometry using BD LSRFortessa.

### Mitochondrial and Cytosol Reactive Oxygen Species Assay using MitoSOX and Dihydroethidium Staining

Drug-treated cells at a concentration of 1.0×10^6^ cells/ml in preheated PBS were incubated with 5 µM mitoSOX Red indicator (Invitrogen, M36008) or 5 µM dihydroethidium (Invitrogen, D11347) at 37°C, 5% CO_2_ for 15 minutes, protected from light. Cells were then washed with preheated PBS, and fluorescent signals generated by mitoSOX or dihydroethidium were detected by flow cytometry using BD LSRFortessa BV605 channel for superoxide detection.(105)

### H460 Xenograft Model in NOD/SCID Mice

The Johns Hopkins University Animal Care and Use Committee approved all animal experiments. Animal care and protocols followed the guidelines of the Institutional Animal Care and Use Committee (IACUC). 1.0 × 10^5^ viable H460 cells in a 1:1 mix of RPMI1640 + 10% FBS and ECM gel (Sigma, E1270) were implanted in the flank of 10-week-old NOD-SCID mice subcutaneously. Drug treatment was randomly assigned to an equal number of male and female mice. Tumor volume was measured using a digital caliper and calculated as 0.5×Length×Width×Height. Drug treatment started at 20 days post-implantation when tumors reached measurable sizes with an average volume of ∼40 mm^3^. Mice were treated with 50 mg/kg MYCi975 (DC chemicals) and 50 mg/kg TMP195 (DC chemicals) following a 5 days-on and 2 days-off schedule and sacrificed when the tumor volume exceeded 2,000 mm^3^. Visualization of the tumor volume data for the MYCi975-treated group stopped on day 25 due to the loss of a mouse. A linear mixed model with random slope fit by the Restricted maximum likelihood (REML) was used to analyze longitudinal tumor volume data for all mouse models. The treatment group:time interaction was used to compare the tumor growth, and the p-value was determined via the Satterthwaite approximation.

### NSCLC Patient-derived Xenograft in NSG Mice

A PDX mouse model (The Jackson Laboratory, TM00302) used in the study was derived from a female patient diagnosed with metastatic lung adenocarcinoma harboring KRAS G12D mutation. 2 mm^3^ fragments of tumor were implanted in the flank of 10-week-old NSG mice. Drug treatment was randomly assigned to an equal number of males and females. Drug treatment was initiated at 25 days post-implantation when tumors reached an average volume of ∼120 mm^3^. Mice were treated with 50 mg/kg MYCi975 (DC chemicals) and 50 mg/kg TMP195 (DC chemicals) following a 5-days-on and 2-days-off schedule and sacrificed when the tumor volume exceeded 2,000 mm^3^.

### Lewis Lung Adenocarcinoma (LLC1) Model in C57BL/6 Mice

1.0 × 10^5^ viable LLC1 cells in 1:1 mix of DMEM + 10% FBS and ECM gel (Sigma, E1270) were implanted on the flank of 10 weeks old C57BL/6 mice subcutaneously. Mice were treated with 50 mg/kg MYCi975 (DC chemicals) and 50 mg/kg TMP195 (DC chemicals) following a 5 days-on and 2 days-off schedule and sacrificed when the tumor volume exceeded 2,000 mm^3^.

### Tumor Sample Collection from LLC1 Model and RNA-seq Library Generation

At 16-day post-treatment, tumors were harvested and processed into a single-cell suspension as described in Newton, J.M., et al.(106) Briefly, the subcutaneous tumor was cut into 1 mm^3^ pieces, followed by incubation with a digestion cocktail (10 mg/mL collagenase I, 2,500 U/mL collagenase IV, 200 U/mL DNase I, 2 mM HEPES in base RPMI-1640 media) for 1 h at 37 °C with shaking at 60 - 100 rpm using an orbital plate shaker. The digestion was stopped by adding a neutralizing buffer (5% FBS, 2 mM EDTA, 2 mM HEPES in base RPMI-1640 media), and the tumor pieces were disaggregated using a 40 µm cell strainer. The cell suspension was centrifuged, and the resulting pellet was incubated with ACK Lysing buffer (Gibco, #A1049201) to lyse red blood cells. The sample was washed and resuspended in a neutralizing buffer, passed through a 40 µm cell strainer, and subsequently enumerated. Tumor cells were enriched using the Miltenyi Tumor Cell Isolation Kit (#130-110-187) using single-cell suspension as input. For Cd11b^+^ population isolation, mononuclear cells were isolated from the single-cell suspension using Lymphoprep (STEMCELL, #07861). Next, the Cd11b^+^ population was isolated using Miltenyi CD11b MicroBeads mouse (#130-126-725). RNA was extracted from the single cell suspension (bulk tumor), isolated Cd11b^+^, and enriched tumor population using the RNeasy Mini Kit (Qiagen, #74104). Following the manufacturer’s protocol, RNA-seq libraries were generated using SMARTer Stranded Total RNA Sample Prep Kit (Takara, 634875). Bioanalyzer and NEBNext Library Quant Kit for Illumina (NEB, E7630) were used to assess the quality and quantity of each library, respectively. These libraries were sequenced on the Illumina Novaseq X Plus instrument (2 x 150bp) for paired-end reads.

### FACS for LLC1 In Vivo Samples

Mononuclear cells isolated from the LLC1 single-cell suspension were stained with Live/dead fixable green dead cell stain (Invitrogen, L23101) and fluorochrome-labeled monoclonal antibodies. The following surface markers were used: NK1.1 (Biolegend, PK136, 108729), CD11b (Biolegend, M1/70, 108724), Gr-1 (Biolegend, RB6-8C5, 108433), CD90.2 (Biolegend, 30-H12, 105335), CD11c (Biolegend, N418, 117333), CD19 (Biolegend, 6D5, 115555), CD69 (Biolegend, H1.2F3, 104543), CD45R/B220 (Biolegend, RA3-6B2, 103207), CD4 (Biolegend, GK1.5, 100455), F4/80 (Biolegend, BM8, 123113), I-A/I-E (Biolegend, M5/114.15.2, 107626), CD8a (BD Bioscience, 53.6.7, 565968), CD45.2 (BD Bioscience, 104, 741092), CD3 (BD Bioscience, 17A2, 741319). Foxp3 / Transcription Factor Staining Buffer Set (eBioScience, 00-5523-00) was used for Foxp3 (MF-14, Biolegend, 126408) staining according to the manufacturer’s protocol. Fluorescent signals were detected by flow cytometry using BD LSRFortessa, and FlowJo_v10.8.1 was used for analysis.

### Transcriptome Analysis

#### CCLE and TCGA RNA-seq Analysis

CCLE RNA-seq read count data from RSEM for the cell lines were acquired from the DepMap portal. The DEseq2 R package(107) was used for differential expression analysis of responders vs. non-responders. Wald statistics derived from the DGE analysis were used as input for GSEA on Hallmark and mitoCarta 3.0 gene sets. RNA-seq read count data from RSEM for the TCGA LUAD samples were acquired from the Broad Firehose portal. Genomic alterations for EGFR and KRAS were defined as putative driver mutations or amplification in these genes. Genomic alterations for STK11 were defined as putative driver mutations or deep deletions in this gene. Genomic alteration data were gathered from the cBioPortal for the TCGA LUAD samples. The DESeq2 R package was used to perform DGE analyses comparing groups harboring distinct genomic alterations. Wald statistics derived from the DGE analysis were used as input for GSEA on Hallmark and mitoCarta 3.0 gene sets. MYC expression levels for the TCGA LUAD samples were derived from the variance stabilized transformation (VST) of RNA-seq counts.

### NSCLC Cell Lines RNA-seq Analysis

Raw FASTQ files were trimmed before downstream analysis using Trimmomatic v0.39, paired-end mode. Processed trimmed files were then aligned using STAR to the Hg38 Homo Sapiens reference genome with GENCODEv38 comprehensive gene annotation. Mapped BAM files generated by STAR were used as input for featureCounts to generate a count matrix with associated annotation. Read count data were subjected to DGE analyses comparing different treatment conditions in each cell line or pooled data using the DEseq2 R package. Gene Set Variation Analysis (GSVA) was performed with VST-normalized expression values on Hallmark or MitoCarta 3.0 gene sets. Differential expression analysis at the pathway level was run with the GSVA output matrices using the limma package.(108) Volcano plots were generated using the EnhancedVolcano package, and the significance of each differentially expressed pathway was determined by FDR-corrected adjusted p-value < 10^−2^. Gene Set Enrichment Analysis (GSEA) on Hallmark cell cycle-related pathways or C3: transcription factor targets gene sets was performed with Wald statistics derived from the DGE analysis with the pooled data from the 4 cell lines or the individual cell line data. A heatmap for the VST-normalized expression of leading-edge genes from suppressed cell cycle-related pathways was generated using the pheatmap package.

### LLC1 Tumor Samples RNA-seq Analysis

Salmon was used for transcript level alignment with raw FASTQ files to gencodevM34 transcript annotation, and the gene-level count matrices were generated using the tximport R package. The DESeq2 package was used for DGE analysis. Gene Set Variation Analysis and Gene Set Enrichment Analysis were performed as described.

### Gene Expression Omnibus Transcriptomic Data Analysis

For RNA-seq data analysis for GSE135800 and GSE246922, raw FASTQ files were downloaded using the SRA toolkit. Salmon was used for transcript level alignment. The DESeq2 package was used for DGE analysis, and Gene Set Enrichment Analysis was performed as described. For microarray data analysis (GSE126453), CEL files were retrieved and loaded into the Oligo R package(109) for RMA algorithm pre-processing of expression data, followed by the DGE using Limma.(108)

### Single-cell RNA-seq Data Visualization

LLC1 single-cell sequencing data with annotated clusters were obtained from the Gene Expression Omnibus (GSE169688, LLC1.1.5Days.Control1, 2, and 3). Seurat R package was used to generate a violin plot representing the expression of Itgam for each cell population in three control LLC1 tumor samples.

## Acknowledgments

Research funding for this work was provided by Van Andel Institute–Stand Up To Cancer Epigenetics Dream Team, Stand Up To Cancer is a division of the Entertainment Industry Foundation (S.B.B.). The Evelyn Grollman Glick Scholar Award (M.J.T.), The Dr. Miriam and Sheldon G. Adelson Medical Research Foundation (S.B.B.), US National Institutes of Health grants: Specialized Program of Research Excellence (SPORE) program, through the National Cancer Institute (NCI), grant P50CA254897 (S.B.B. and M.J.T.), and P30CA006973 (SKCCC Core Grant). The content is solely the responsibility of the authors and does not necessarily represent the official views of the NIH. J.P. was supported by the Kwanjeong Educational Foundation and Margaret Lee fellowship. Sequencing experiments were conducted at the Genetic Resources Core Facility (RRID #: SCR_018669), Johns Hopkins Department of Genetic Medicine, Baltimore, MD.

## Author Contributions

Conceptualization: JP, MJT

Resources: RCY

Methodology: JP, MJT, and YC

Investigation: JP, MJT, YC, and JJC

Visualization: JP

Funding acquisition: SBB, MJT

Supervision: SBB, MJT

Writing – original draft: JP, MJT

Writing – review & editing: JP, MJT, YC, JA, JDO, and SBB

## Conflicts of Interest

S.B.B. is on the Scientific Advisory Board for Mirati Therapeutics Inc. S.B.B. is a consultant for MDxHealth. S.B.B. is an inventor of MSP, which is licensed to MDxHealth in agreement with Johns Hopkins University (JHU). S.B.B. and JHU are entitled to royalty sales shares.

## Supplementary Figure Legends

**Fig. S1. MYCi975 plus TMP269 reduces cell viability in NSCLC cell lines. A,** Volcano plot of GSE135800 RNA-seq. Blue dots represent differentially expressed genes with adjusted p-value < 0.05 and |log_2_ fold-change| > log_2_1.4 between MYCi975 (8 µM, 24 h) and vehicle-treated prostate cancer PC3 cells. Each dot represents one gene. **B,** Volcano plot of GSE126453 microarray generated with pooled data from three NSCLC cell lines (A549, H1299, H1975). Blue dots represent differentially expressed genes with adjusted p-value < 0.05 and |log_2_ fold-change| > log_2_1.4 between Omomyc (12.8 µM, 72 h) and vehicle-treated NSCLC cells. Reanalysis of array data. Black dots represent class IIa HDAC genes, including HDAC1/2/3/6/8/10/11 (A and B). **C** and **D,** Dose-response matrices for percent viability in responsive cell lines **(C)** or non-responsive cell lines **(D)**. Values are reported as means from two (H520, A549, H661) or three (H1792, H23, H1299, H1755, H596, H226, HCC4006) independent experiments. Each cell line was treated with combination drugs as indicated for 72 h. **E** and **F,** MYCi975 dose-response curves for normalized proliferation in the absence or presence of TMP269 (0, 10, 25 µM) in responsive cell lines **(E)** or non-responsive cell lines **(F)**. **G** and **H,** The color on the dose-response surfaces represents the synergy score derived from the Bliss model in responsive cell lines **(G)** or non-responsive cell lines **(H)**. Cool color indicates synergy and warm color indicates antagonism.

**Fig. S2. MYCi975 plus TMP195 reduces cell viability in NSCLC cell lines. A,** Dose-response matrices for percent viability. Values are reported as means from three (H1703) or four (H460, H838, H1755, HCC827, H441) independent experiments. Each cell line was treated with combination drugs as indicated for 72 h. **B,** MYCi975 dose-response curves for normalized proliferation in the presence or absence of TMP195 (0, 10, 25 µM). **C,** The color on the dose-response surfaces represents the synergy score derived from the Bliss model. Cool color indicates synergy and warm color indicates antagonism. **D,** HDAC-Glo Class IIa Assay on cells treated with indicated drugs for 24 h. Mean ± SD. n=2 biological replicates. p-values defined as *<0.05, **< 0.01, ***<0.001, ****<0.0001; were calculated using one-way ANOVA followed by Tukey’s multiple comparisons test.

**Fig. S3. Molecular features of NSCLC cell lines derived from the CCLE dataset and human NSCLC tumors derived from the TCGA dataset. A,** Principal Component Analysis (PCA) plot depicting PC1 and PC2 for 18 cell lines in the screening panel. The drug response and subtype of each cell line are annotated. **B,** Volcano plot of CCLE RNA-seq data. Orange dots represent differentially expressed genes with adjusted p-value < 0.05 and |log_2_ fold-change| > log_2_1.4 between Responsive vs. nonresponsive cell lines. Teal dots represent genes with adjusted p-value ≥ 0.05 and |log_2_ fold-change| > log_2_1.4. Black dots represent genes with adjusted p-value ≥ 0.05 and |log_2_ fold-change| ≤ log_2_1.4. Each dot represents one gene. Cancer Cell Line Encyclopedia (CCLE) RNA-seq data from the DepMap Portal was used for the analysis (A and B). **C,** Normalized protein levels for HDAC4, HDAC5, HDAC7, and HDAC9. Cell lines with missing values were excluded. CCLE proteomics data was obtained through the DepMap Portal. The Mann-Whitney test derived the p-values. R: responders, NR: non-responders (A, B, and C). **D−F,** MYC RNA expression in TCGA LUAD samples without or with EGFR (n=398 vs. n=55) **(D)**, KRAS (n=302 vs. n=151) **(E)**, or STK11 (n=401 vs. n=52) **(F)** genetic alterations. p-values were derived by the Mann-Whitney test. **G** and **H,** Oncoprint plot representing STK11, KRAS, or EGFR genetic alterations in TCGA LUSC **(G)** or LUAD **(H)** samples. Each box represents a single patient with indicated genomic alterations. **I** and **J,** Principal Component Analysis plots. TCGA RNA-seq data obtained from the Broad Firehose portal was used for the analysis. Genomic alteration status for EGFR, KRAS, and STK11 are color-coded. **K,** A schematic diagram representing the mutational and transcriptional profiles associated with combination drug responses.

**Fig. S4. Hallmark pathways and genes involved in cell cycle progression are suppressed by the combination treatment. A,** GSEA with cell cycle-related Hallmark pathways based on differential expression analysis using pooled data from 4 responsive cell lines (H460, H838, H1703, H1755) treated with Combo vs. Vehicle. Running enrichment scores and the distribution of gene sets in the ranked list of genes are depicted. **B,** Heatmap representing the expression of leading-edge genes in suppressed cell cycle-related pathways derived from GSEA. Z-score scaled vst-normalized expression data is depicted. Yellow indicates a positive Z-score, and purple indicates a negative Z-score.

**Fig. S5. Combination treatment induces stronger transcriptional response upon combination treatment in responders compared to non-responders. A,** MYC protein levels were detected by western blotting in H460, H1703, HCC827, and H441 cells treated with MYCi975 at indicated doses for 48 h. All immunoblots are representative of two biological replicates. **B,** H460 cells were transfected with an empty vector or MYC overexpression vector, and the protein was collected. MYC protein levels were detected by western blotting. **C,** Volcano plots from the RNA-seq data (48 h treatment) from 2 responsive cell lines (H460 and H1703) and 2 non-responsive cell lines (HCC827 and H441) for Combo vs. Vehicle. Orange dots represent differentially expressed genes with adjusted p-value < 0.05; teal dots represent genes with adjusted p-value ≥ 0.05. Each dot represents one gene. n=2 biological replicates. **D,** RNA levels for AURKA, AURKB, PLK1, PLK4, CDC25A, and E2F2 were measured by RT-qPCR in H460, H1703, HCC827, and H441 cells treated with indicated agents for 48 h. Mean ± SD. n=3 biological replicates. p-values defined as *<0.05, **< 0.01, ***<0.001 were calculated using one-way ANOVA followed by Tukey’s multiple comparisons test for indicated condition vs. vehicle with log2(fold change over vehicle) data.

**Fig. S6. ATAC-seq analysis of drug-treated H838 cells. A,** Volcano plots of ATAC-seq consensus peaks. Orange dots represent differentially accessible regions with adjusted p-value < 0.05. Each dot represents one peak. H838 cells were treated with vehicle, MYCi975 (5 µM), TMP195 (25 µM), or both for 24 h. n=4 biological replicates. **B,** Heatmaps representing ATAC-seq peaks within differentially accessible regions (Combo vs. Vehicle) for indicated samples in H838 cells. **C,** Genomic annotation of Differentially Accessible Regions (DAR) for indicated comparisons in H460 and H838 cells. **D,** GREAT (the Genomic Regions Enrichment of Annotations Tool) analysis on cell-cycle related pathways in H838 cells. **E,** Lists of transcription factors (TF) with motifs that are positively or negatively enriched in the combo group compared to the vehicle group in ATAC-seq data and with their target gene sets enriched in RNA-seq in H838. **F,** Running enrichment scores and the distribution of gene sets in the ranked list of genes for transcription factor target gene sets identified by integrating RNA- and ATAC-seq data for H838 cells.

**Fig. S7. MYCi plus class IIa HDACi reduces tumor growth in vivo. A** and **B,** Mean volumes of H460 xenograft tumors implanted in NOD/SCID mice treated with the drugs as indicated **(A)**. Graphing of mean volumes for the MYCi975 group stopped at day 25 due to a loss of mouse. Spider plots depict individual tumor growth **(B)**. n=7. **C** and **D,** Mean volumes of patient-derived xenografts (LUAD) harboring KRAS G12D mutation implanted in NSG mice treated with the drugs as indicated **(C)**. Spider plots depict individual tumor growth **(D)**. n=5 for vehicle, TMP195, Combo. n=6 for MYCi975. Treatment schema below each plot: black indicates on-treatment, and white indicates off-treatment (A and C). Mean ± SEM. (A and C) Linear mixed model by the Restricted maximum likelihood (REML) approach was used to derive p-values (A and C). **E–G,** % weight changes of H460 xenografts **(E)**, patient-derived xenografts **(F)**, or LLC1 **(G)** implanted mice treated with indicated drugs.

**Fig. S8. RNA-seq analysis of LLC1 bulk tumor samples. A,** Volcano plots from differential expression analysis with the LLC1 bulk tumor RNA-seq for Combo, TMP195, or MYCi975 vs. Vehicle. Orange dots represent differentially expressed genes with adjusted p-value < 0.1. **B,** GSEA with Hallmark gene sets based on differential expression analysis with bulk tumor from TMP195 or MYCi975 vs. Vehicle-treated mice. n=3 for TMP195. n=4 for MYCi975 and vehicle.

**Fig. S9. High-level immune profiling of LLC1 tumor and RNA-seq analysis of LLC1 enriched tumor samples. A,** Gating Strategy for immune profiling from LLC1 tumor-bearing mice. **B,** The percentage of immune cells in total Cd45^+^ cells is derived from flow cytometry analysis as described in **(A)**. n=5 for vehicle and TMP195, n=3 for MYCi975, and n=4 for Combo. B cells (Cd3^−^B220^+^Cd11c^−^), Foxp3^−^Cd4^+^ T cells (B220^−^Cd3^+^Cd4^+^Foxp3^−^), Foxp3^+^Cd4^+^ T cells (B220^−^Cd3^+^Cd4^+^Foxp3^+^), Cd69^+^Foxp3^−^Cd4^+^ T cells (B220^−^Cd3^+^Cd4^+^Cd69^+^Foxp3^−^), Cd8^+^ T cells (B220^−^Cd3^+^Cd8^+^), Cd69^+^Cd8^+^ T cells (B220^−^Cd3^+^Cd8^+^Cd69^+^), Macrophages (B220^−^Cd3^−^F4/80^+^), MDSCs (myeloid-derived suppressor cells) + Neutrophils (B220^−^Cd3^−^F4/80^−^Nk1.1^−^Gr1^+^Cd11b^+^), DCs (dendritic cells) (B220^−^Cd3^−^F4/80^−^Nk1.1^−^Cd11c^+^MHCII^+^), pDCs (plasmacytoid dendritic cell) (Cd3^−^B220^+^Cd11c^+^), and NK (natural killer) cells (B220^−^Cd3^−^Nk1.1^+^) are presented. **C,** Volcano plots from differential expression analysis with the LLC1 enriched tumor cell RNA-seq for Combo, TMP195, or MYCi975 vs. Vehicle. Orange dots represent differentially expressed genes with adjusted p-value < 0.1. **D,** GSEA with Hallmark gene sets based on differential expression analysis with enriched tumor cells from TMP195 or MYCi975 vs. Vehicle-treated mice. **E,** GSEA with MitoCarta 3.0 gene sets based on differential expression analysis with enriched tumor cells from TMP195 vs. Vehicle-treated mice. MYCi975 treatment resulted in no significant enrichment of MitoCarta 3.0 gene sets. n=4 (C−E). **F** and **G,** GSEA with Hallmark **(F)** or MitoCarta 3.0 **(G)** gene sets based on differential expression analysis with Cd45^−^ tumor cells derived from ICB- relapsed vs. parental LLC (GSE246922). n=3.

**Fig. S10. RNA-seq analysis of LLC1 tumor-infiltrative Cd11b+ population. A,** Single cell sequencing data for live Cd45^+^ cells sorted from dissected LLC tumor cell suspension obtained from GSE169688 (LLC1.1.5Days.Control1, 2, and 3). Violin plot represents the expression of Itgam (A gene name for Cd11b) for each cell population defined in Thumkeo, Dean, et al.(76) **B,** Volcano plots from differential expression analysis with the isolated Cd11b^+^ RNA-seq for Combo, TMP195, or MYCi975 vs. Vehicle. Orange dots represent differentially expressed genes with adjusted p-value < 0.1. n=4 for vehicle, MYCi975, and Combo, n=3 for TMP195. **C,** GSEA with Hallmark gene sets based on differential expression analysis with isolated Cd11b^+^ RNA-seq comparing Combo, TMP195, or MYCi975 vs. Vehicle-treated mice. **D** and **E,** GSEA with GOBP positive regulation of vasculature development **(D)** and GOBP establishment of endothelial barrier **(E)** based on differential expression analysis with isolated Cd11b^+^ RNA-seq. Running enrichment scores and the distribution of gene sets in the ranked list of genes are depicted. **F,** GSEA with cell type signature gene sets based on differential expression analysis with bulk tumor from Combo or MYCi975 vs. Vehicle-treated mice. n=3 for Combo. n=4 for MYCi975 and vehicle. TMP195 treatment did not significantly enrich endothelial cell-associated cell type signature gene sets. **G,** Heatmap representing the expression of endothelial cell marker genes in bulk tumor treated with indicated drugs. Z-score scaled vst-normalized expression data. Yellow indicates a positive Z-score, and purple indicates a negative Z-score.

