## Supplementary Figures and Tables for "MYC plus class IIa HDAC inhibition potentiates mitochondrial dysfunction in non-small cell lung cancer"

### Supplementary Figure 1

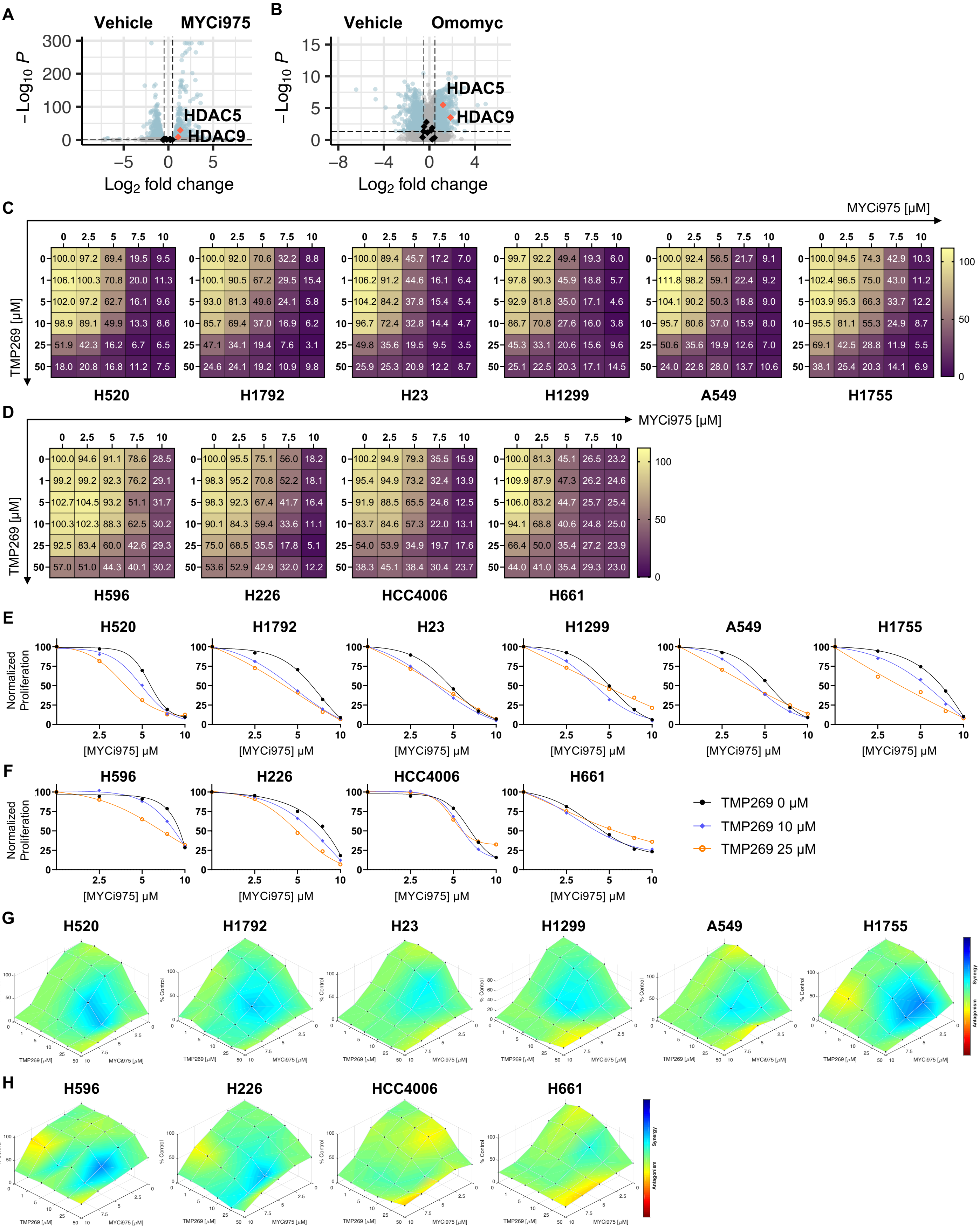

Supplementary Figure 2

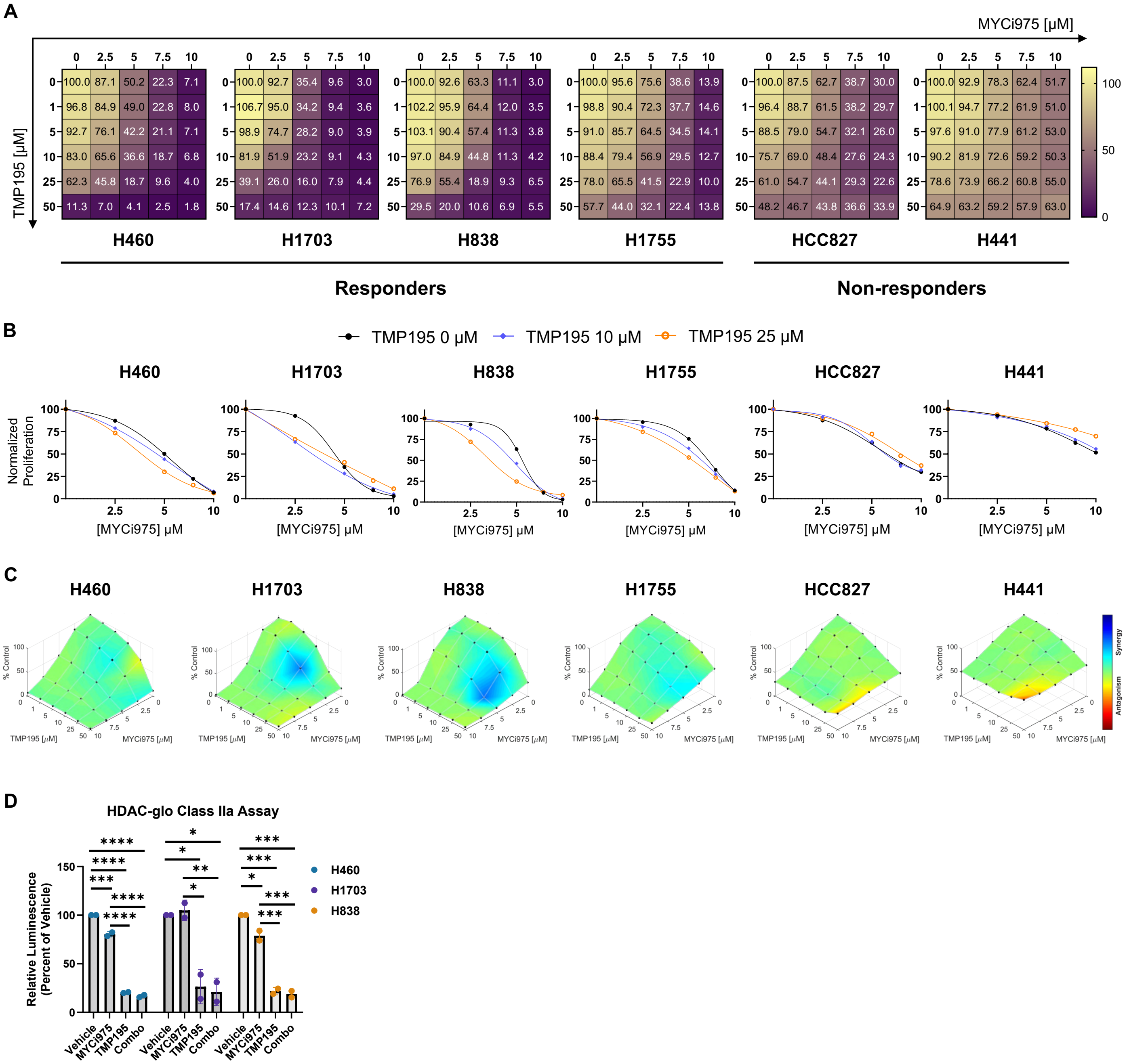

### Supplementary Figure 3

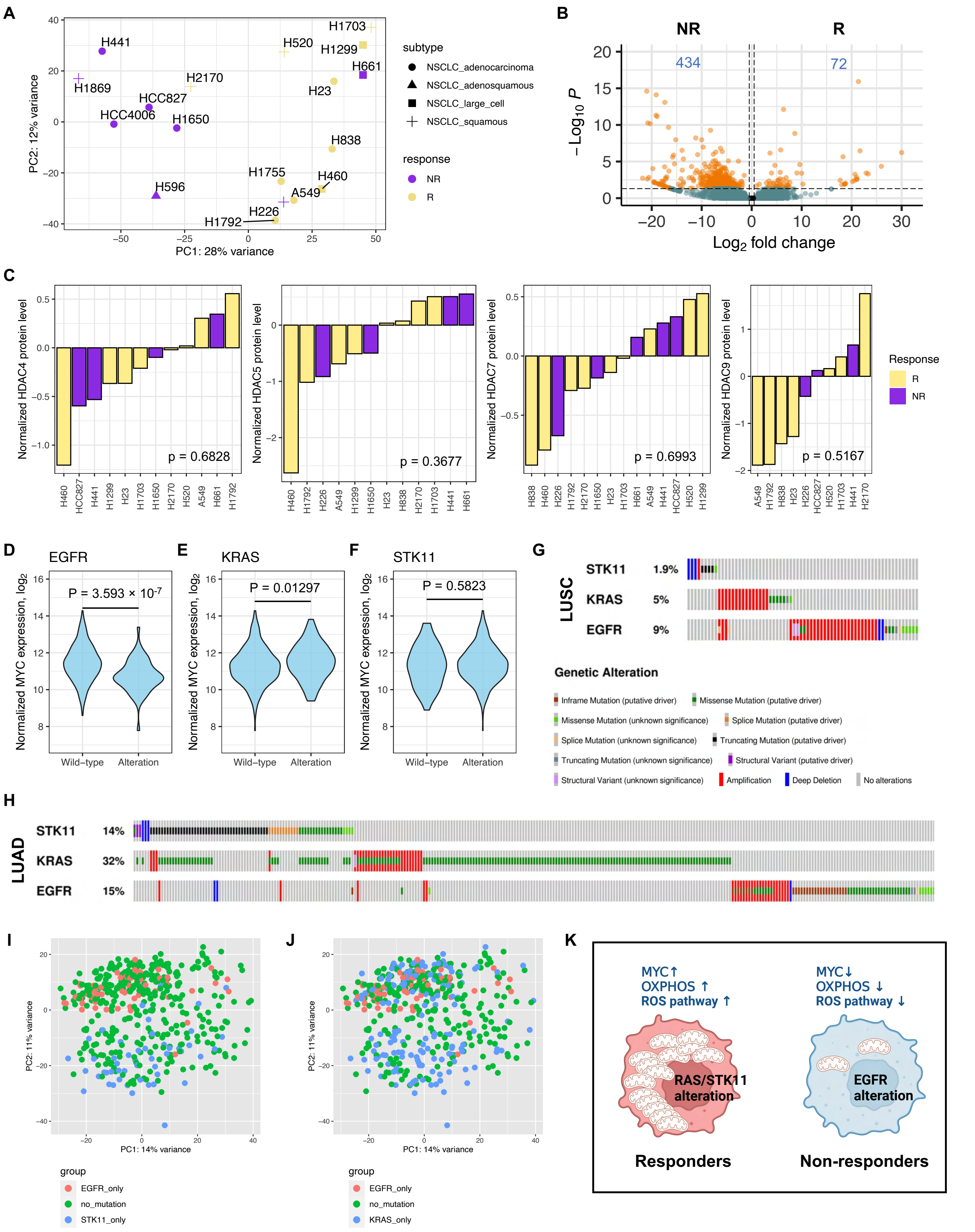

Supplementary Figure 4

A

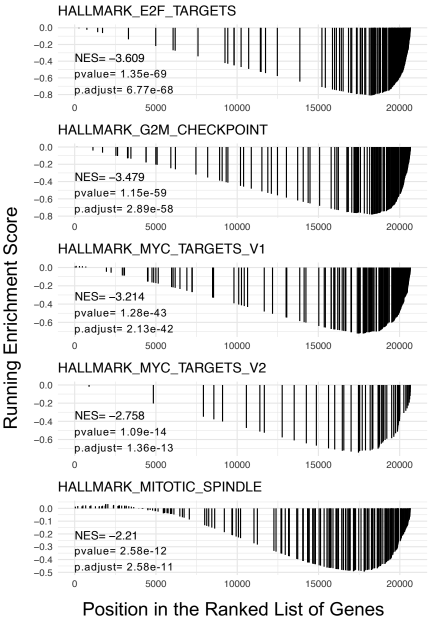


B

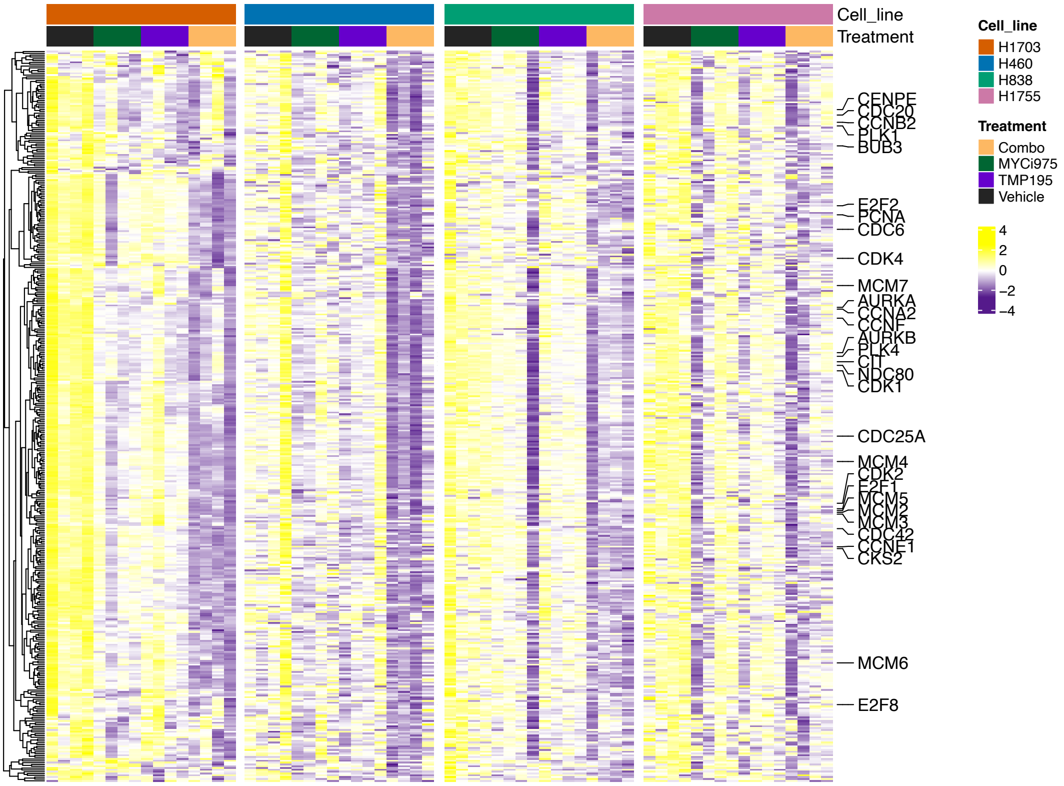

Supplementary Figure 5

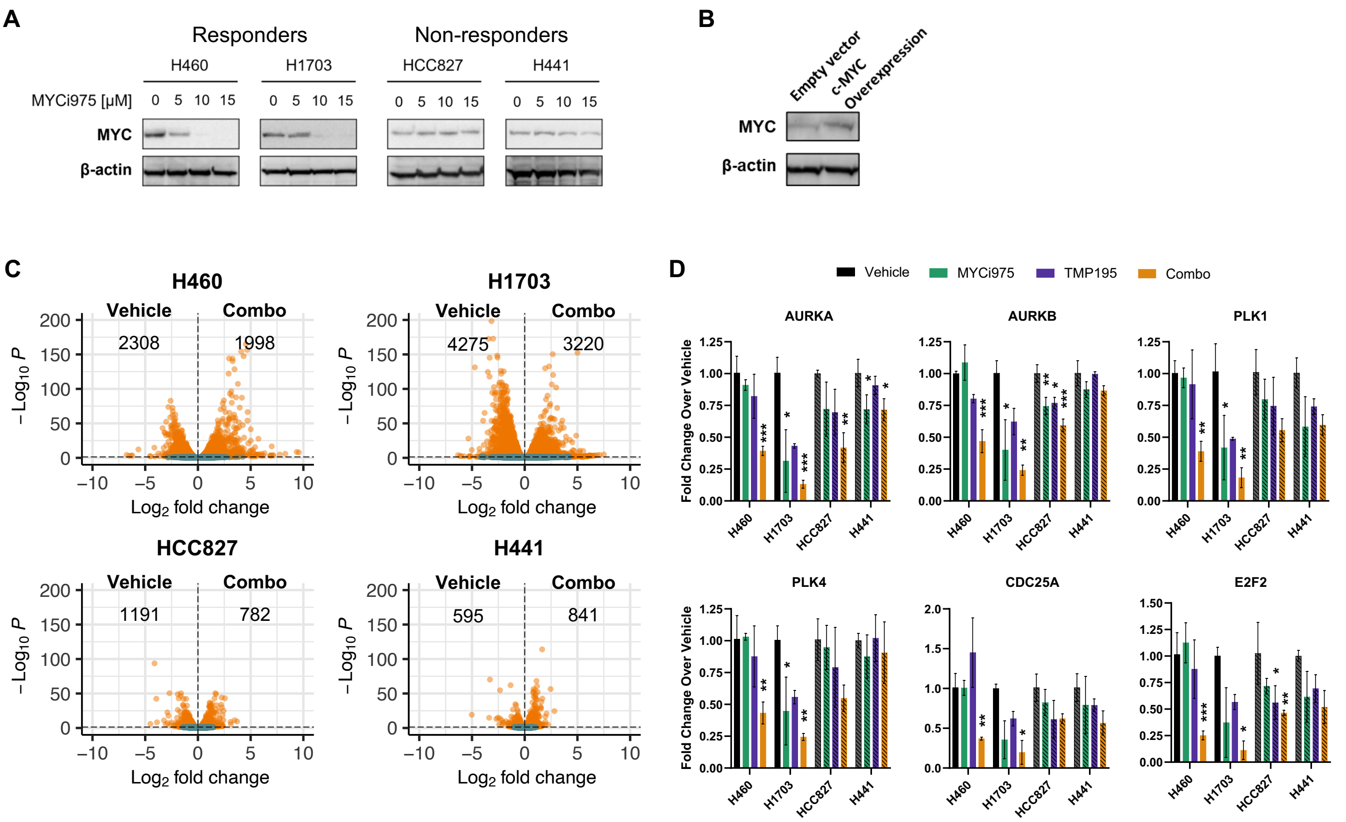

Supplementary Figure 6

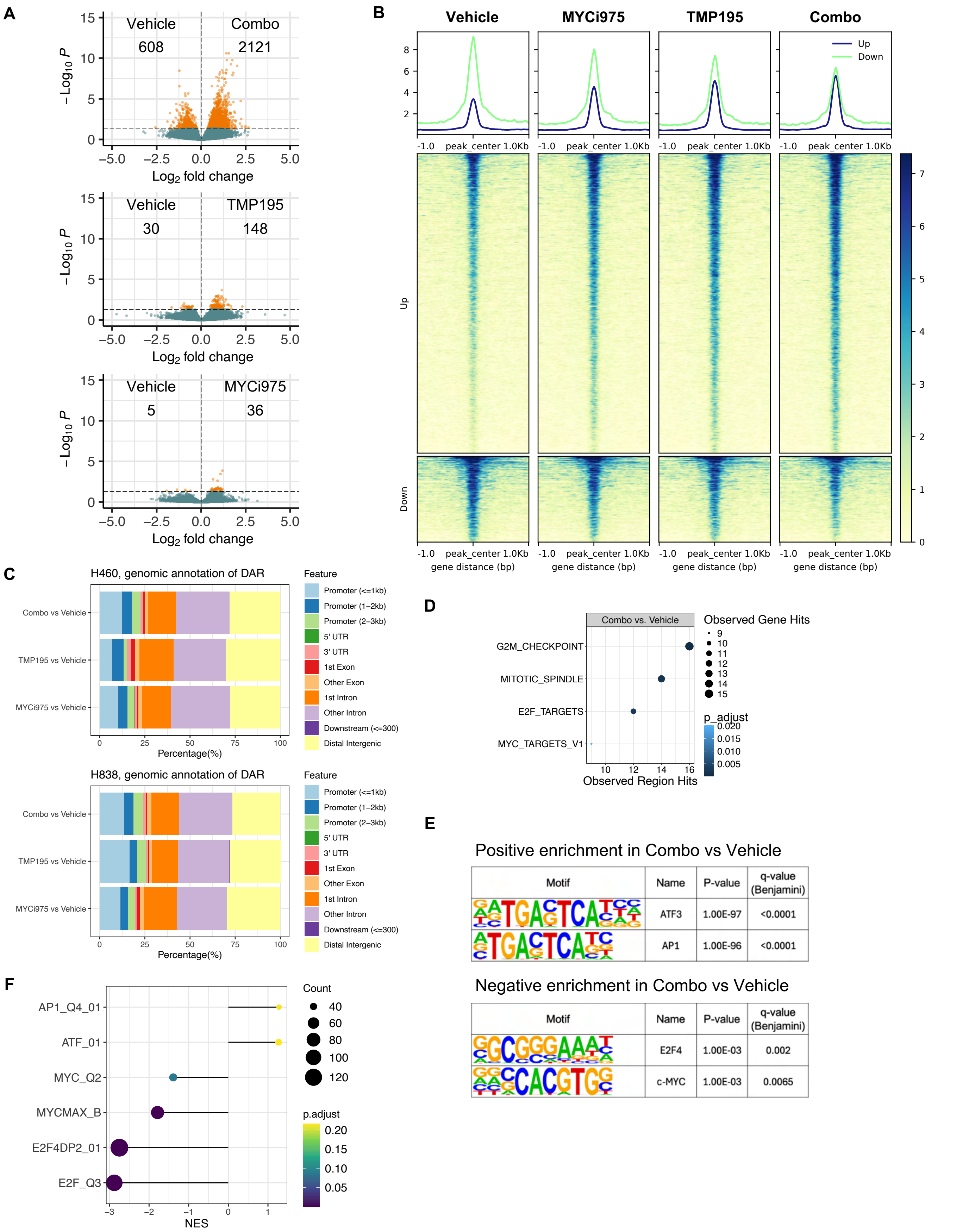

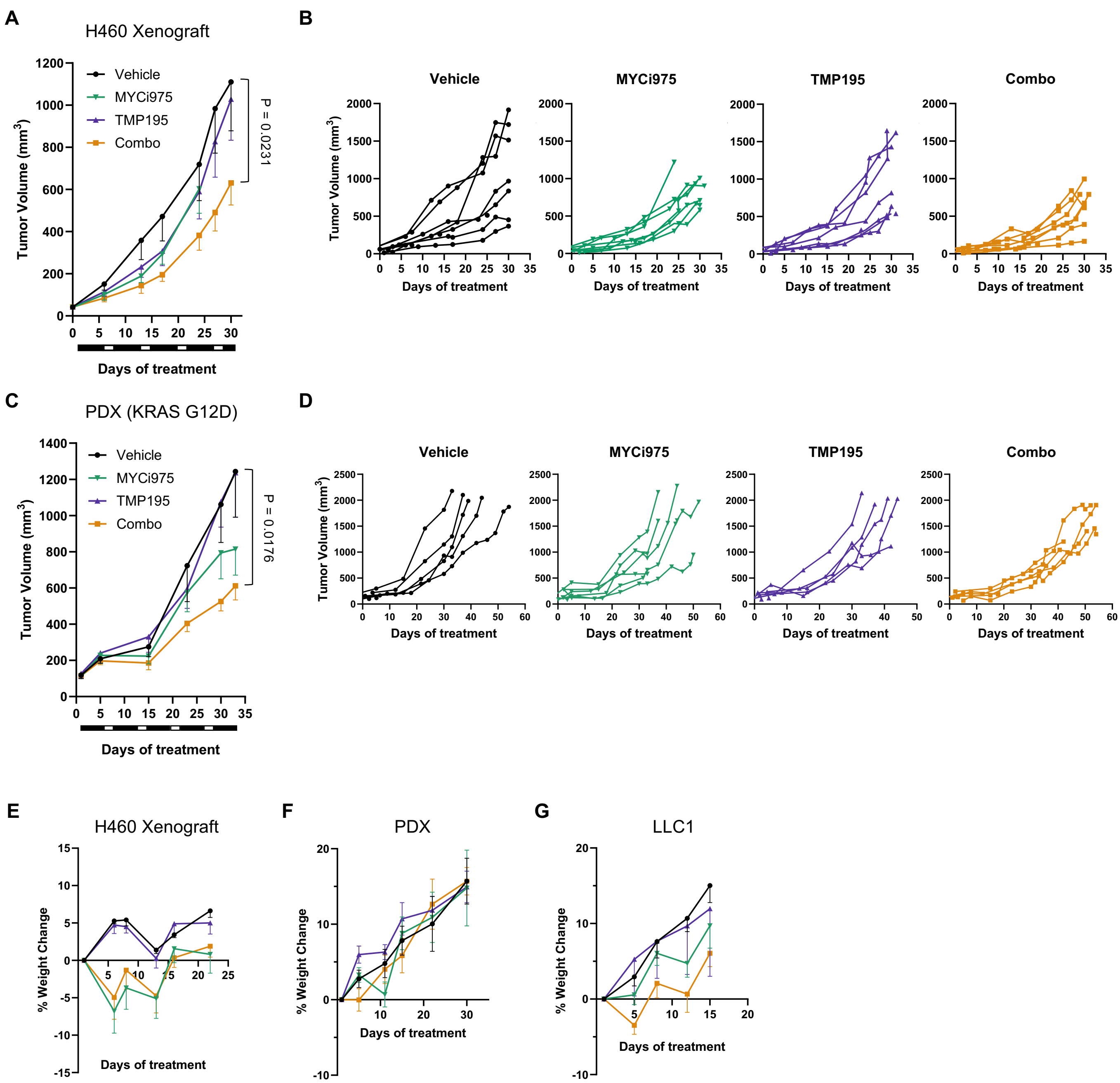
A
H460 Xenograft
Vehicle
MYCi975
TMP195
Combo
P = 0.0231
Tumor Volume (mm³)
Days of treatment
B
Vehicle
MYCi975
TMP195
Combo
Tumor Volume (mm³)
Days of treatment
C
PDX (KRAS G12D)
Vehicle
MYCi975
TMP195
Combo
P = 0.0176
Tumor Volume (mm³)
Days of treatment
D
Vehicle
MYCi975
TMP195
Combo
Tumor Volume (mm³)
Days of treatment
E
H460 Xenograft
% Weight Change
Days of treatment
F
PDX
% Weight Change
Days of treatment
G
LLC1
% Weight Change
Days of treatment

Supplementary Figure 8

A

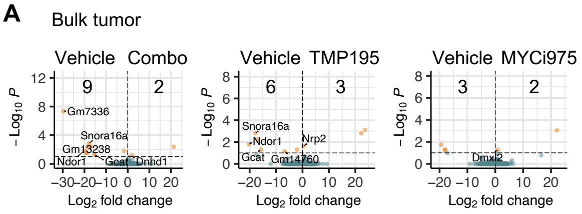


B

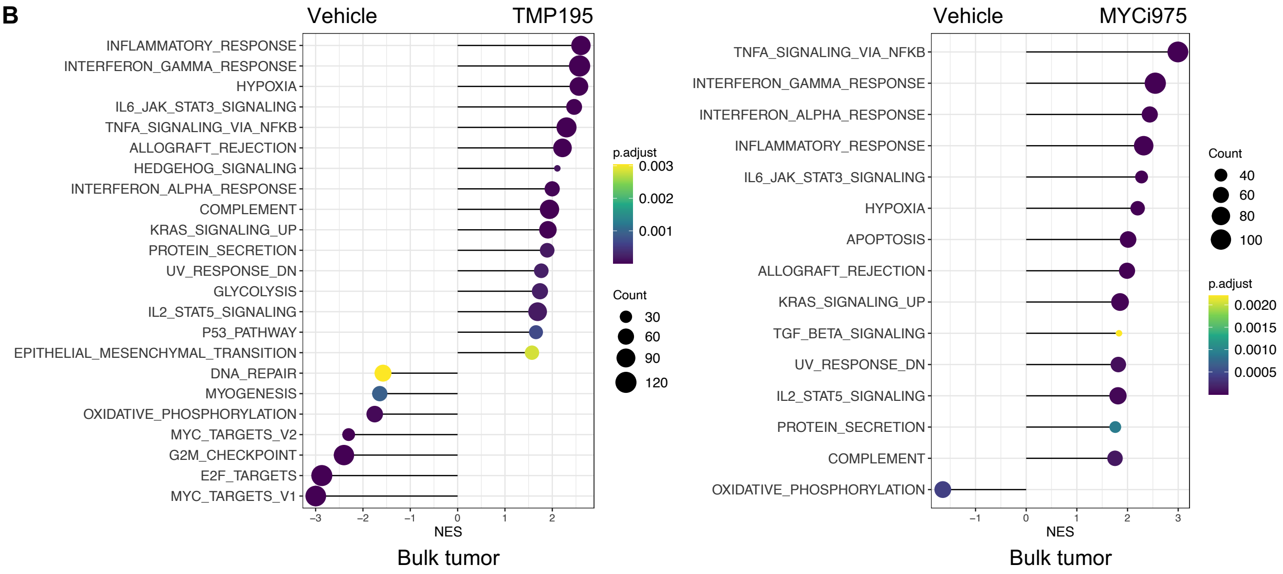

### Supplementary Figure 9

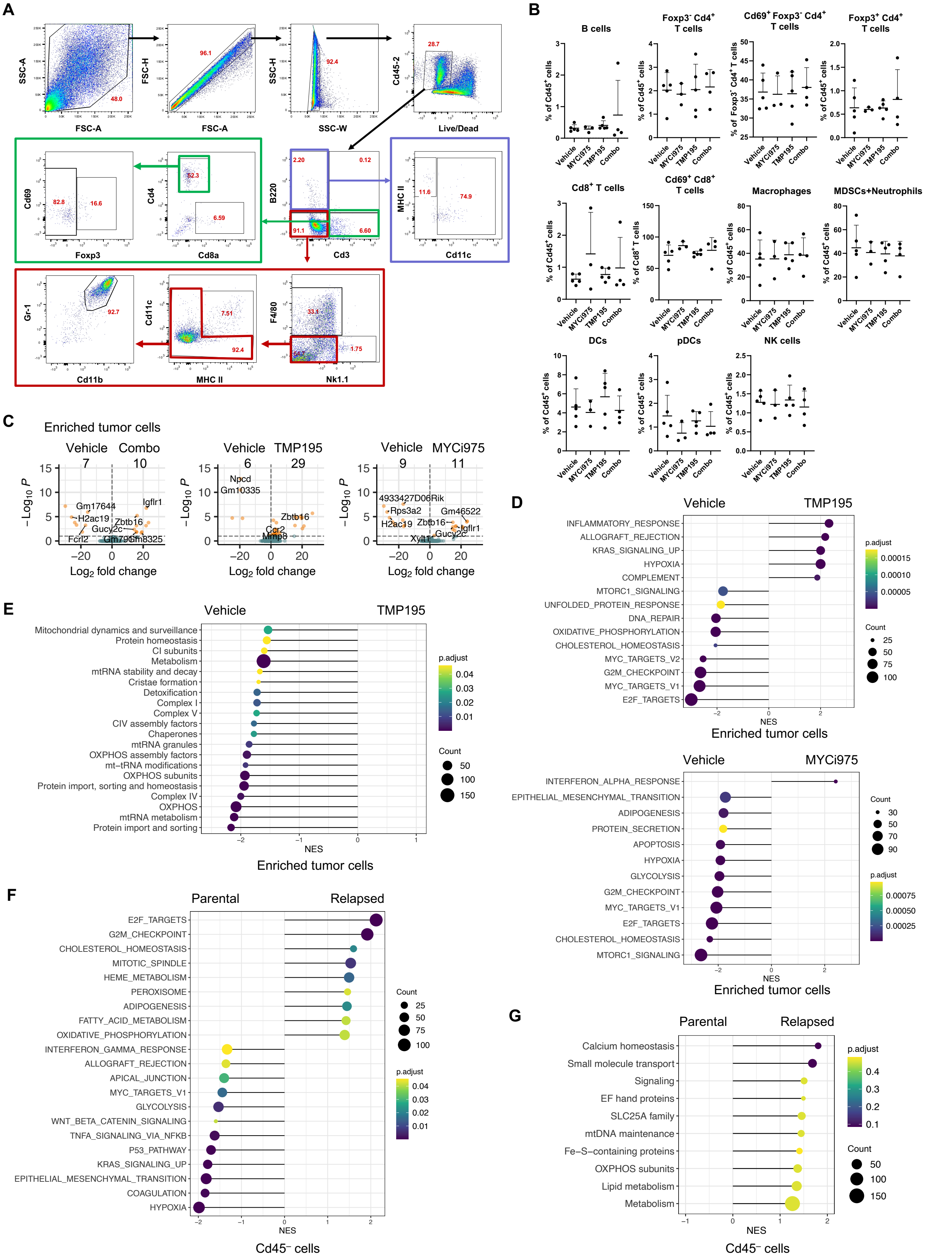

Supplementary Figure 10

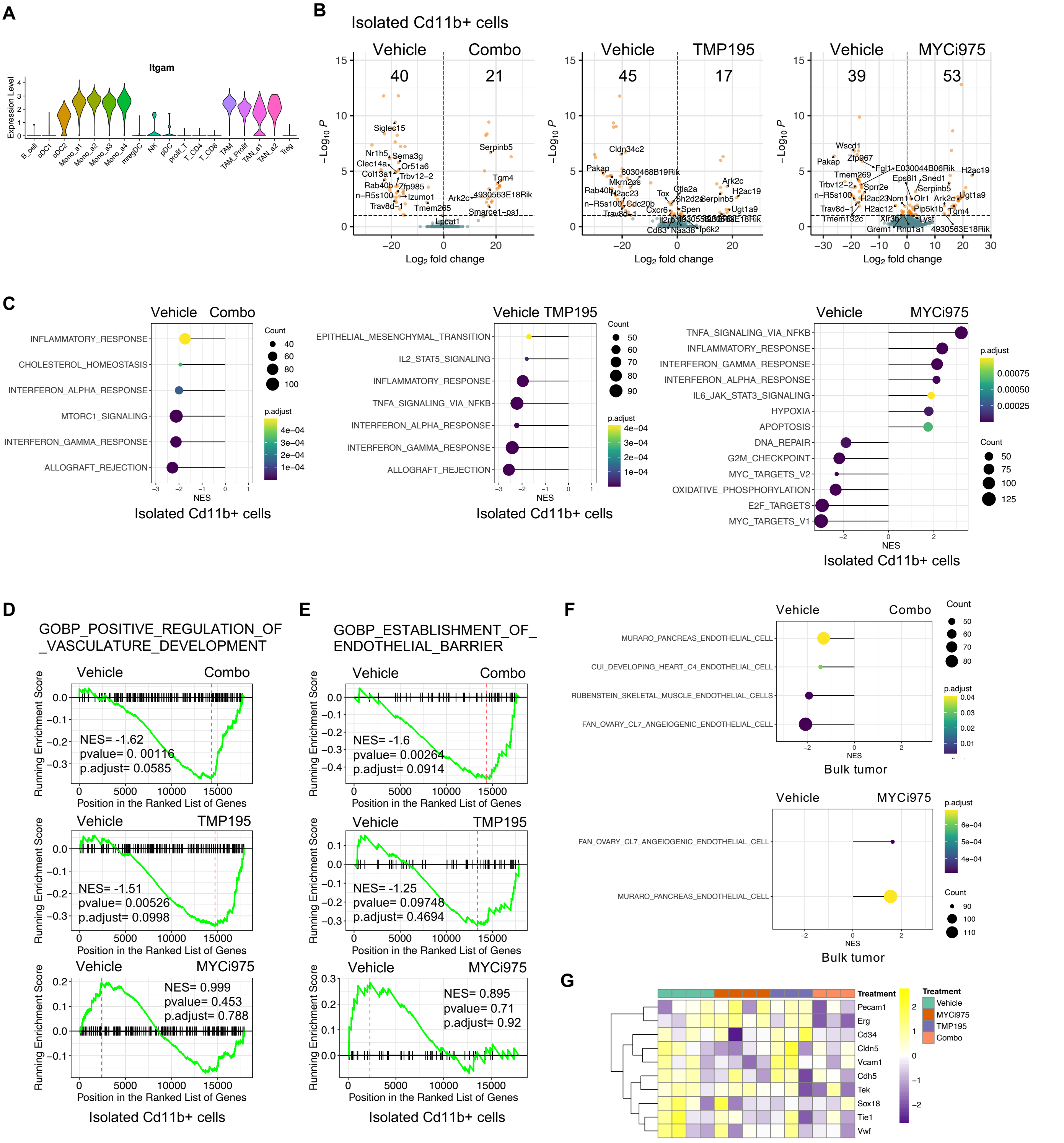

| Gene | Genomic Alteration | R (+/-) | NR (+/-) | p-value | p.adjust |
|---|---|---|---|---|---|
| **CDKN2A** | Hotspot mutation | 3/7 | 1/7 | 0.5882 | 0.70584 |
| **EGFR** | Hotspot mutation | 0/10 | 3/5 | 0.06863 | 0.20589 |
| **KRAS** | Hotspot mutation | 4/6 | 1/7 | 0.3137 | 0.6274 |
| **PIK3CA** | Hotspot mutation | 1/9 | 1/7 | 1 | 1 |
| **STK11** | Hotspot mutation | 5/5 | 0/8 | 0.03595 | 0.20589 |
| **TP53** | Hotspot mutation | 7/3 | 7/1 | 0.5882 | 0.70584 |
| **CDKN2A** | Deletion | 4/6 | 2/6 | 0.638 | 0.7975 |
| **KRAS** | Amplification | 3/7 | 1/7 | 0.5882 | 0.7975 |
| **EGFR** | Amplification | 0/10 | 3/5 | 0.06863 | 0.6863 |
| **NKX2-1** | Amplification | 1/9 | 2/6 | 0.5588 | 0.7975 |
| **FOXA1** | Amplification | 2/8 | 1/7 | 1 | 1 |
| **BCL2L1** | Amplification | 1/9 | 1/7 | 1 | 1 |
| **TERT** | Amplification | 1/9 | 2/6 | 0.5588 | 0.7975 |
| **NSD3** | Amplification | 2/8 | 0/8 | 0.4771 | 0.7975 |
| **FGFR1** | Amplification | 2/8 | 0/8 | 0.4771 | 0.7975 |
| **MYC** | Amplification | 3/7 | 1/7 | 0.5882 | 0.7975 |
| **CDKN2A** | Hotspot mutation/Deletion | 7/3 | 3/5 | 0.3416 | 0.3416 |
| **EGFR** | Hotspot mutation/Amplification | 0/10 | 4/4 | 0.02288 | 0.06864 |
| **KRAS** | Hotspot mutation/Amplification | 6/4 | 1/7 | 0.06561 | 0.098415 |

**Supplementary Table 1. List of cancer driver genes altered in 18 NSCLC cell lines**

Related to Fig. 2E. The number of responsive or non-responsive cell lines with or without each genomic alteration is shown. P-values were derived from Fisher’s exact test for genomic alteration vs. response. FDR-adjusted p-values were calculated using the Benjamini–Hochberg method.

| Gene | Forward Primer | Reverse Primer |
|---|---|---|
| AURKA | GTCCCACCTTCGGCATCCTA | TCGAATGACAGTAAGACAGGGC |
| AURKB | GGCTCAAGGGAGAGCTGAAG | ACATTGTCTTCCTCCTCAGGG |
| PLK1 | GGCACAGTTTCGAGGTGGAT | CCACGGGGTTGATGTGCTTG |
| PLK4 | AGCGGCGGTTTAGAGAGC | TTTCCAACTTTAAAATCCTCGATCT |
| CDC25A | CAGGGTCTGGGCAGTGATTA | GTCCAATGGCCCAGGAGAAT |
| E2F2 | CTGAGCTTCAAGCACCTGACT | ATCTGCAGGTTGTCCTCAGTCC |
| MYC | CCCTACCCTCTCAACGACAG | TTCTTGTTCCTCCTCAGAGTCG |
| TBP | TGTGCTCACCCACCAACAA | CTGCTCTGACTTTAGCACCTGTT |
| UBC | CGTCGCAGCCGGGATTTG | CACGAAGATCTGCATTGTCAAG |

**Supplementary Table 2. List of forward and reverse primers for genes tested in RT-qPCR experiments**
